# Expanding the known nucleorhabdovirus world: the final chapter in a trilogy exploring the hidden diversity of plant-associated rhabdoviruses

**DOI:** 10.1101/2025.05.09.653113

**Authors:** Nicolás Bejerman, Ralf Georg Dietzgen, Humberto Debat

## Abstract

Nucleorhabdoviruses, classified into four distinct genera within the family *Rhabdoviridae* (subfamily *Betarhabdovirinae*), are plant-infecting viruses, characterized by nucleus-associated, enveloped, bacilliform virions. Nucleorhabdoviruses possess an unsegmented, single-stranded, negative-sense RNA genome (ca. 12-15.2 kb) that encodes six to eight proteins. Here, by exploring large publicly available metatranscriptomics datasets, we report the identification and genomic characterization of 47 novel viruses with genetic and evolutionary hallmarks of nucleorhabdoviruses. These viruses are associated with 45 distinct host plant species and were previously hidden in public database. Our findings significantly broaden the known host range of nucleorhabdoviruses, including the first nucleorhabdovirus associated with ferns and the first gammanucleorhabdoviruses linked to dicot hosts. Genetic divergence and evolutionary analyses suggest that all these novel viruses likely represent members of novel species. Phylogenetic reconstruction indicates that ten novel viruses are related to alphanucleorhabdoviruses, 25 to betanucleorhabdoviruses, eight to deltanucleorhabdoviruses and four to gammanucleorhabdoviruses. This study constitutes the final chapter in a trilogy chronicling a data mining expedition into the cryptic diversity of plant-associated rhabdoviruses, a journey that began with varicosaviruses and continued with cytorhabdoviruses. These new findings yield the most comprehensive phylogeny of nucleorhabdoviruses to date, significantly expanding our genomic understanding and illuminating the phylogenetic relationships and evolutionary dynamics of this virus group. Further, this work underscores the value of large-scale sequence data mining in advancing our understanding of the hidden world of plant rhabdoviruses.

## 1. Introduction

In the current era of metagenomics, the rapid discovery of novel viruses has revealed a vast and diverse evolutionary landscape of replicating entities, presenting significant challenges in their systematic classification (Koonin et al., 2021). In response to this complexity, various strategies have emerged, culminating in a comprehensive proposal for establishing a “megataxonomy” of the viral world (Koonin et al., 2024). Despite extensive efforts to characterize the viral component of the biosphere, it is evident that only a tiny fraction, likely less than one percent, of viral diversity has been thoroughly explored (Geoghegan and Holmes, 2017; Dominguez-Huerta et al., 2023; Hou et al., 2024; Lu et al., 2025). As a result, our understanding of the global virome remains limited, particularly regarding its immense diversity and the interactions between viruses and their hosts (Dolja et al., 2020; Edgar et al., 2022; Koonin et al., 2022; Mifsud et al., 2022). To address this gap, researchers have increasingly turned to mining publicly available transcriptome datasets derived from High-Throughput Sequencing (HTS), which offers a rapid and cost-effective approach to viral discovery (Edgar et al., 2022; Bejerman et al., 2023; Charon et al., 2024; Sidharthan and Baranwal, 2024). This data-driven strategy has proven particularly valuable given the sheer volume of freely available datasets within the Sequence Read Archive (SRA), maintained by the National Center for Biotechnology Information (NCBI), which continues to expand at an extraordinary pace. Although the data in SRA represent a partial and potentially biased sample of Earth’s biodiversity, the NCBI-SRA remains an exceptionally efficient and economical resource for the identification of novel viruses (Lauber and Seitz, 2022). The tool ‘Serratus’ (Edgar et al., 2022) has proven invaluable, enabling large-scale data mining and accelerating viral sequence discovery at an unprecedented pace. In the realm of virus taxonomy, a growing consensus has emphasized the necessity of incorporating viruses identified solely through metagenomic data into the official classification framework of the International Committee on Taxonomy of Viruses (ICTV) (Simmonds et al., 2017). This shift underscores the vital role of metagenomic approaches in expanding our understanding of the global virome and modernizing taxonomic systems to accommodate the ever-growing known diversity of viruses (Simmonds et al., 2023).

The family *Rhabdoviridae* comprises viruses with negative-sense single-stranded RNA genomes that infect a wide range of hosts, including plants, amphibians, fish, mammals, reptiles, insects, and other arthropods (Dietzgen et al., 2017; Walker et al., 2022). While most rhabdovirus genomes are unsegmented, recent discoveries have revealed plant-associated rhabdoviruses with bi-segmented and trisegmented genomes (Bejerman et al., 2021; Bejerman et al., 2022; Walker et al., 2022; Bejerman et al., 2023). Historically, *Nucleorhabdovirus* was a single genus of plant-associated rhabdoviruses, characterized by replication within the nuclei of infected plant cells. This genus was recently divided into three genera *Alphanucleorhabdovirus, Betanucleorhabdovirus* and *Gammanucleorhabdovirus* within the subfamily *Betarhabdovirinae* (Dietzgen et al., 2020; Walker et al., 2022). These genera currently include 17 (https://ictv.global/report/chapter/rhabdoviridae/rhabdoviridae/alphanucleorhabdovirus), 26 (https://ictv.global/report/chapter/rhabdoviridae/rhabdoviridae/betanucleorhabdovirus) and two species (https://ictv.global/report/chapter/rhabdoviridae/rhabdoviridae/gammanucleorhabdovirus), respectively. In 2024, a fourth genus, *Deltanucleorhabdovirus*, was created to accommodate two novel species that are phylogenetically distinct from members of the other three genera (Simmonds et al., 2024). All nucleorhabdoviruses share a conserved genome organization, typically encoding six conserved canonical genes in the order 3′-nucleocapsid protein (N)-phosphoprotein (P)–movement protein (P3)-matrix protein (M)-glycoprotein (G)–large polymerase (L)-5′. Additionally, an accessory gene of unknown function has been identified in 13 out of the 45 described nucleorhabdoviruses (nine alphanucleorhabdoviruses, two betanucleorhabdoviruses and both known gammanucleorhabdoviruses), while only one member has two accessory genes (Walker et al., 2022). The coding regions are separated by conserved gene junction sequences and are flanked by 3′ leader and 5′ trailer sequences, which have partially complementary ends that may form a panhandle structure during replication (Dietzgen et al., 2017).

This study represents the final chapter in a trilogy documenting a data mining expedition into the cryptic diversity of plant-associated rhabdoviruses. This trilogy started in 2022 with the exploration of varicosaviruses (Bejerman et al., 2022), followed in 2023, by a focus on cytorhabdoviruses (Bejerman et al., 2023). A prequel to this trilogy was published in 2021, centered on the exploration of plant-associated rhabdoviruses using the Transcriptome Shotgun Assembly (TSA) database (Bejerman et al., 2021). In this third installment, through mining of publicly available sequence data, we identified 47 novel nucleorhabdoviruses associated with plants. These discoveries not only expand the known host range of nucleorhabdoviruses but also double the number of described members, providing fresh insights into their evolutionary landscape.

## 2. Material and Methods

### 2.1 Identification of nucleorhabdovirus-like sequences from public plant RNA-seq datasets

We analyzed the Serratus database using the Serratus Explorer tool (Edgar et al., 2022), employing as query sequences the predicted RNA-dependent RNA polymerase protein (RdRP) of nucleorhabdoviruses available at the NCBI Refseq database. The SRA libraries showing matches to the query sequences (alignment identity > 45%; score > 10) were selected for further analysis.

### 2.2 Sequence assembly and virus identification

Virus discovery was done as previously described (Debat et al., 2023; Bejerman et al., 2023). Briefly, raw nucleotide sequence reads from each SRA experiment that matched the query sequences in the Serratus platform were downloaded from their corresponding NCBI BioProjects (**Tables 1-4**). Datasets were preprocessed by trimming and filtering with the Trimmomatic v0.40 tool as implemented in http://www.usadellab.org/cms/?page=trimmomatic with standard parameters except quality requirement was raised from 20 to 30 (initial ILLUMINACLIP step, sliding window trimming, average quality required = 30). The filtered reads were assembled *de novo* using rnaSPAdes using standard parameters on the Galaxy server (https://usegalaxy.org/). The resulting transcripts obtained from *de novo* transcriptome assembly were subjected to bulk local BLASTX searches (E-value < 1e^-5^) against nucleorhabdovirus Refseq protein sequences available at https://www.ncbi.nlm.nih.gov/protein?term=txid11306 [Organism]. Viral sequence hits were examined in detail. Tentative virus-like contigs were curated (extended and/or validated) through iterative mapping of filtered reads from each SRA library. In each round, a subset of reads corresponding to the contig of interest was extracted and used to extend the sequence, which was then used as a new query for the next iteration. This process was repeated until no further extension was possible. The final extended and polished transcripts were reassembled using Geneious v8.1.9 (Biomatters Ltd.) alignment tool with high sensitivity parameters.

### 2.3 Bioinformatics tools and analyses

#### 2.3.1 Sequence analyses

Open reading frames (ORFs) were predicted with ORFfinder (minimal ORF length of 120 nt, genetic code 1, https://www.ncbi.nlm.nih.gov/orffinder/). The functional domains and overall architecture of the translated gene products were analyzed using InterPro (https://www.ebi.ac.uk/interpro/search/sequence-search) and the NCBI Conserved domain database -CDD v3.20 (https://www.ncbi.nlm.nih.gov/Structure/cdd/wrpsb.cgi) with an e-value threshold of 0.01. To further annotate divergent predicted proteins, HHPred and HHBlits as implemented in https://toolkit.tuebingen.mpg.de/#/tools/were employed. Transmembrane domains were predicted using the TMHMM version 2.0 tool (http://www.cbs.dtu.dk/services/TMHMM/), while signal peptides were predicted using the SignalP version 6.0 tool (https://services.healthtech.dtu.dk/services/SignalP-6.0/ ). The predicted proteins were then subjected to NCBI-BLASTP searches against the non-redundant protein sequences (nr) database to identify potential homologs.

#### 2.3.2 Pairwise sequence identity

Percentage nucleotide (nt) sequence identities of the coding-region, as well as the amino acid (aa) sequence identities of the predicted L proteins, were calculated for all viruses identified in this study, as well as those available in the NCBI database, using SDTv1.2 (Muhire et al., 2014) based on multiple sequence alignments generated with MAFFT v7.505 (https://mafft.cbrc.jp/alignment/software) with standard parameters. Virus names and abbreviations of nucleorhabdoviruses already reported are shown in **Supplementary Table 1**.

#### 2.3.3 Phylogenetic analysis

Phylogenetic analyses based on the predicted L protein of all plant nucleorhabdoviruses, listed in **Table S1**, were done using MAFFT v7.505 https://mafft.cbrc.jp/alignment/software, with multiple aa sequence alignments using FFT-NS-i as the best-fit model. The aligned aa sequences were used as the input in MEGA11 software (Tamura et al., 2021) to generate phylogenetic trees by the maximum-likelihood method (best-fit model = WAG + G + F). Local support values were computed using bootstraps with 1,000 replicates. L proteins of selected betacytorhabdoviruses were used as outgroup.

## 3. Results and Discussion

### 3.1. Summary of discovered viral sequences

It is highly probable that the majority of nucleorhabdoviruses do not induce overt symptoms in their hosts, a phenomenon previously reported for varicosaviruses (Bejerman et al., 2022) and cytorhabdoviruses (Bejerman et al., 2023). This hypothesis is supported by several recent high-throughput sequencing (HTS) studies in which nucleorhabdoviruses were identified in apparently asymptomatic plants (Rivarez et al., 2023; Safarova et al., 2024; Reyes-Proaño et al., 2023; Hu et al., 2023; Chen et al., 2022C; Li et al., 2022B; Medberry et al., 2022; Belete et al., 2022; Zhou et al., 2020; Bhat et al., 2020; Baek et al., 2019; Cao et al., 2019; Gaafar et al., 2019). In addition, eight nucleorhabdoviruses were previously discovered through mining publicly available transcriptome datasets from *Bacoppa monnieri* and *Lotus corniculatus* (Sidharthan and Baranwal, 2021; Debat and Bejerman 2019) as well as through analysis of the metatranscriptomic data housed in the TSA database (Bejerman et al., 2021). These findings strongly suggest that a substantial number of additional nucleorhabdoviruses remain to be discovered through comprehensive mining of publicly available metatranscriptomic datasets. The SRA database likely harbors a substantial number of hidden nucleorhabdovirus sequences, particularly in datasets derived from asymptomatic plants, where viral presence is not typically anticipated. Despite its immense potential, the SRA remains underexplored in this context. The advent of the powerful Serratus platform (Edgar et al., 2022) has revolutionized the mining of this resource, enabling rapid and large-scale screening that would otherwise be prohibitively labor-intensive and time-consuming (Bejerman et al., 2023; Charon et al., 2024). Leveraging this tool, we conducted the most extensive search to date on nucleorhabdovirus RdRP sequences. This effort led to the identification and full coding regions assembly of 47 novel nucleorhabdoviruses, effectively doubling the number of known viruses in this group. Moreover, our findings significantly broaden the known host range, including the first nucleorhabdovirus associated to ferns, and the first gammanucleorhabdoviruses associated to dicot hosts. In contrast to the cytorhabdoviruses, which display diverse genome organizations with many members having unique genome architectures and some of them lacking some genes thought to be conserved among plant rhabdoviruses (Bejerman et al., 2021, 2023), the nucleorhabdoviruses as a group display a relatively conserved genomic architecture. All these viruses possess the canonical six genes present in plant rhabdoviruses, and some of them have only one additional ORF, and only one has two additional ORFs. This genomic conservatism suggests a more homogeneous evolutionary trajectory, where only limited gene acquisitions have occurred. These observations support the hypothesis that these viruses evolved from viruses of plant-feeding arthropods that acquired movement proteins and assorted RNAi suppressors through recombination with preexisting plant viruses (Dolja et al., 2020). Their relatively constrained evolution may be attributed to a tight ecological association with arthropod vectors, leading to a conservative genomic path (Whitfield et al., 2018), in stark contrast to the more evolutionarily promiscuous cytorhabdoviruses (Bejerman et al., 2021, 2023).

### 3.2 Alphanucleorhabdovirus

The full-length coding regions of ten novel alphanucleorhabdoviruses were assembled in this study (**Table 1**), increasing the number of members of this genus by 0.6-fold. These viruses were associated with ten different plant hosts (**Table 1**), including six herbaceous dicots, one woody dicot, and three gramineous monocots. This host range mirrors that of previously reported alphanucleorhabdoviruses, which include herbaceous dicots (7/17), woody dicots (2/17) and monocots (7/17), with one member lacking a defined host (https://ictv.global/report/chapter/rhabdoviridae/rhabdoviridae/alphanucleorhabdovirus). These findings suggest that alphanucleorhabdoviruses have followed a complex host adaptation trajectory, with different lineages evolving to infect herbaceous dicots, woody dicots, or monocots. This adaptation is likely influenced by the host preferences of their insect vectors. Among plant rhabdoviruses, there is a strong correlation between phylogenetic relationships and vector type (Dietzgen et al., 2020), and alphanucleorhabdoviruses have been reported to be transmitted by either planthoppers or leafhoppers (Dietzgen et al., 2020). Therefore, it is plausible to speculate that these novel alphanucleorhabdoviruses are also transmitted by leafhopper or planthopper vectors. However, further studies should be carried out to identify and confirm their vectors.

The genomic organization of the ten alphanucleorhabdoviruses identified in this study is notably diverse. Seven of these viruses lack additional accessory genes and display the canonical 3′-N-P–P3-M-G-L-5′ (**Table 1, Fig. 2b**), consistent with the seven previously reported alphanucleorhabdoviruses (https://ictv.global/report/chapter/rhabdoviridae/rhabdoviridae/alphanucleorhabdovirus) (**Fig. 2B**). Two of the novel alphanucleorhabdovirusess have an accessory ORF between the G and L genes displaying a 3′-N-P–P3-M-G-P6-L-5′ genomic organization, also reported in two already known members of the genus (**Fig. 2B**). The remaining novel virus has an accessory ORF between the N and P genes displaying a 3′-N-X-P–P3-M-G-L-5′ genomic organization, which is similar to seven previously reported alphanucleorhabdoviruses (**Fig. 2B**). The P6 protein encoded by rice yellow stunt virus has been proposed to function as a suppressor of RNA silencing (Guo et al., 2013); however, no conserved functional domain has been identified in the P6 proteins of either the viruses described here or those of rice yellow stunt virus (Huang et al., 2003) and wheat yellow striate virus (Liu et al., 2018). As such, the putative function of this additional gene is unknown. Interestingly, P6 appears to be exclusive to monocot-infecting alphanucleorhabdoviruses. Future research should focus on the functional characterization of this elusive protein to determine the potential evolutionary advantage conferred by its acquisition. On the other hand, the putative function of X protein has been more clearly elucidated. Recent work demonstrated that X is a non-virion protein required for efficient plant infection and symptom induction (Wang et al., 2024).

The consensus gene junction sequences of the ten novel alphanucleorhabdoviruses identified in this study are highly conserved, both among themselves, and compared with those of previously reported alphanucleorhabdoviruses (**Table 5**), supporting a shared evolutionary history within the genus. Pairwise nt sequence identity values across the complete-coding genome of the ten novel viruses and those from known alphanucleorhabdoviruses ranged between 56.6% and 73.9% (**Table S2**). These moderate levels of divergence suggest that the extent of uncharacterized “viral dark matter” within this genus may be limited.

The phylogenetic analysis based on the L protein aa sequence showed that the ten novel viruses grouped with 17 known alphanucleorhabdoviruses in a well-supported clade (**Fig. 1**), thus, indicating a common evolutionary history. Within this distinct group of 27 viruses, several subclades can be observed (**Fig.2A**), often grouping viruses with similar genome architectures and host types. Although some exceptions exist, such as Agave tequilana virus 1 and cassava virus 1 (**Fig. 2A**), this pattern suggests a coevolutionary trajectory between alphanucleorhabdoviruses and their plant hosts. Interestingly, among monocot-infecting viruses, several have acquired an accessory gene between the G and L genes, while several herbaceous dicot-infecting viruses tend to harbor an additional gene between the N and P genes. One virus with an unassigned host also features an accessory gene between the N and P genes, and another one between the M and G genes (**Fig. 2B**).

**Figure 1.**
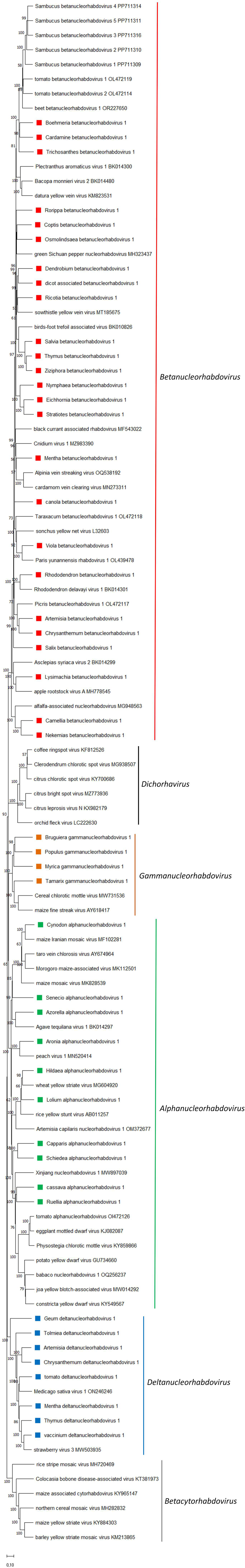
Maximum-likelihood phylogenetic tree based on amino acid sequence alignments of the complete L gene of all nucleorhabdoviruses reported so far and in this study constructed with the WAG + G + F model. The scale bar indicates the number of substitutions per site. Bootstrap values following 1,000 replicates are given at the nodes, but only the values above 50% are shown. The Alpha-, Beta-, Gamma- and Deltanucleorhabdoviruses identified in this study are noted with green, red, orange, and blue rectangles, respectively. Betacytorhabdoviruses were used as outgroup.

**Figure 2.**
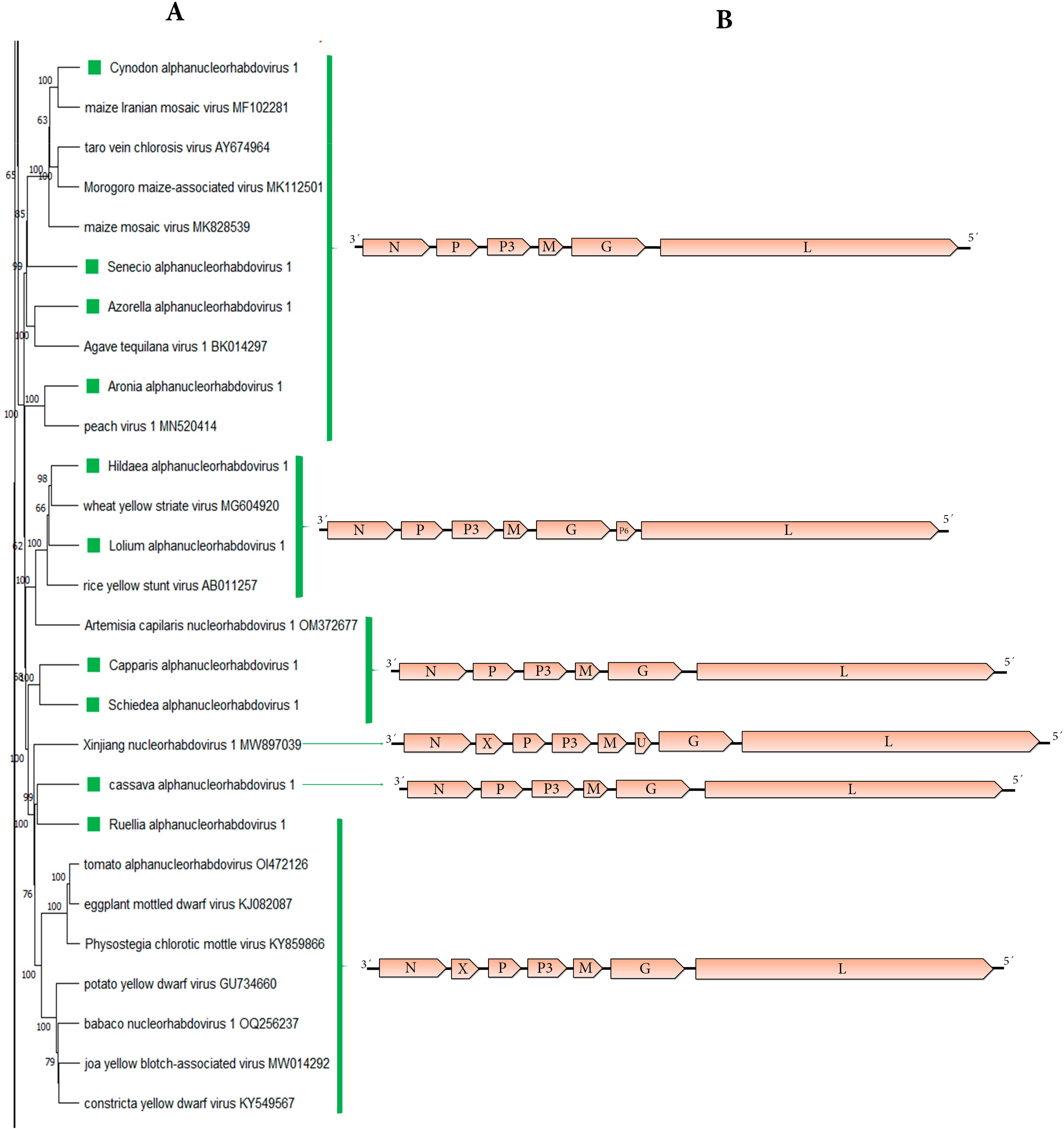
A: The maximum-likelihood phylogenetic tree shown in Fig. 1 was cropped to show only those viruses included in the genus *Alphanucleorhabdovirus*. The viruses identified in this study are noted with green rectangles. B: genomic organization of the viral sequences used in the phylogeny.

According to current species demarcation criteria, viruses assigned to different species within the genus should share less than 75% nucleotide identity across their full genome sequence and occupy different ecological niches, as evidenced by differences in host species and/or arthropod vectors associations (Walker et al., 2022). Therefore, the ten novel alphanucleorhabdoviruses identified in this study should be classified as new species in the genus *Alphanucleorhabdovirus*.

### 3.3 Betanucleorhabdovirus

The full-length coding regions of 25 novel betanucleorhabdoviruses were assembled in this study (**Table 2**), effectively doubling the number of proposed members within this genus. These viruses were associated with 24 known plant hosts and one unassigned host (**Table 2**). Of the identified host plants, eighteen were herbaceous dicots, two were woody dicots, three were monocots, and one was a fern (**Table 2**). This is in line with reported hosts from known betanucleorhabdoviruses which are either herbaceous dicots (18/26), or woody dicots (7/26) (https://ictv.global/report/chapter/rhabdoviridae/rhabdoviridae/betanucleorhabdovirus). Recently, the first betanucleorhabdovirus associated with a monocot host was identified in *Alpinia* (Larrea-Sarmiento et al., 2024). The three monocot-associated novel viruses reported here were found in hosts from distinct monocot families (**Table 2**). Interestingly, no virus was associated with a member of the family *Poaceae*; which are the common monocot host of alpha- and gammanucleorhabdoviruses. Moreover, we expanded the host range of nucleorhabdoviruses identifying the first nucleorhabdovirus associated with a fern. To date, the only other group of plant rhabdoviruses reported in ferns are members of the *Varicosavirus* genus (Bejerman et al., 2022).

Among all plant rhabdoviruses studied so far, there is a strong correlation between phylogenetic relationships and vector types (Dietzgen et al., 2020). Many members of the *Betanucleorhabdovirus* genus are known to be transmitted by aphids (Dietzgen et al., 2020). We therefore predict that the novel betanucleorhabdoviruses reported here are likely aphid-transmitted. Betanucleorhabdoviruses exhibit a complex pattern of host adaptation. While most members are adapted to infect herbaceous dicots, some to woody dicots, others to monocots, and now one to ferns. This evolutionary diversification is likely influenced by the host range and feeding preferences of their aphid vectors. Although aphids can colonize both dicots and monocots (Doring, 2014), it has been suggested that they feed more successfully on dicots (Van Bell and Will, 2016), which may partially explain the predominance of dicot-associated betanucleorhabdoviruses.

The genomic organization of the 25 betanucleorhabdoviruses identified in this study is conserved, as only two distinct genomic structures were observed (**Table 2**; **Fig. 3B**). Twenty-two viruses lack additional accessory genes and have a genome with the canonical order 3′-N-P–P3-M-G-L-5′ (**Table 1, Fig. 2b**), which is similar to most previously reported betanucleorhabdoviruses (24/26) (**Fig. 3B**); while the other three novel betanucleorhabdoviruses have an accessory ORF between the M and G genes displaying a 3′-N-P–P3-M-U-G-L-5′ genomic organization, similar to two previously reported betanucleorhabdoviruses (**Fig. 3B**). Therefore, few betanucleorhabdoviruses acquired an additional ORF during their evolution. No conserved functional domain was identified in this additional protein, named “U”, in those viruses identified in this study, as well as in the U protein encoded by apple rootstock virus A (Baek et al., 2019) and alfalfa-associated nucleorhabdovirus (Gaafar et al., 2019). Therefore, the function of this additional gene is unknown. Future studies should focus on the functional characterization of the U proteins to gain critical insights into their biological roles and to elucidate the potential selective advantage conferred by their acquisition.

**Figure 3.**
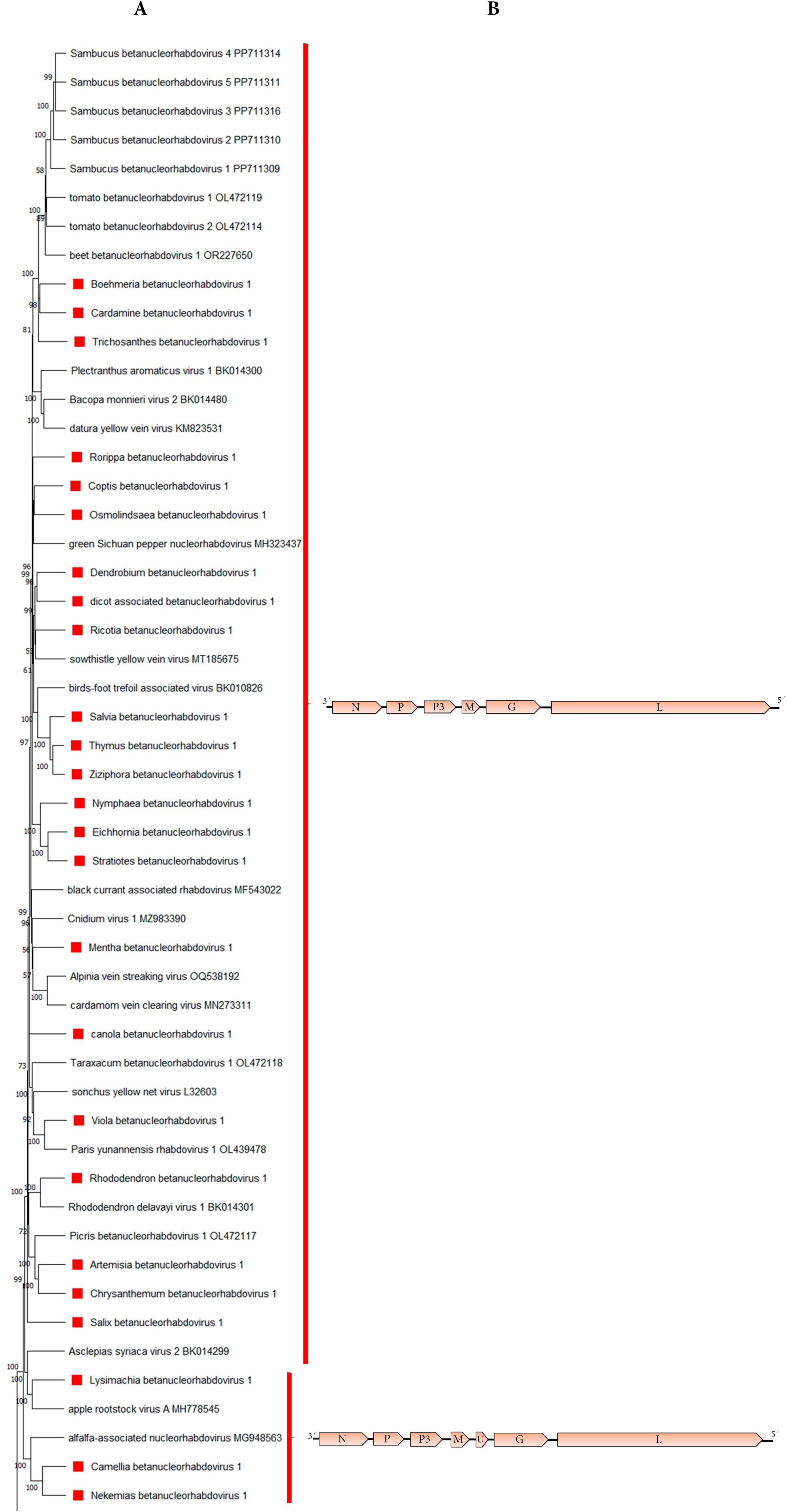
A: The maximum-likelihood phylogenetic tree shown in Fig. 1 was cropped to show only those viruses included in the genus *Betanucleorhabdovirus*. The viruses identified in this study are noted with red rectangles. B: genomic organization of the viral sequences used in the phylogeny.

The consensus gene junction sequences of the novel betanucleorhabdoviruses identified in our study are identical among themselves and to those of previously reported betanucleorhabdoviruses (**Table 5**), supporting a shared evolutionary history for these viruses.

Pairwise nt sequence identity values between the complete-coding genome of the 25 novel viruses and those from known betanucleorhabdoviruses do not vary significantly, ranging between 55.5% and 74.9% (**Table S3**); suggesting that there is limited uncharacterized diversity within betanucleorhabdoviruses. The phylogenetic analysis based on the L protein aa sequence showed that the 25 novel viruses grouped with 26 known betanucleorhabdoviruses in a well-supported, monophyletic clade (**Fig. 1**), further supporting their shared ancestry. Within this distinct group of 51 viruses, there are two major subclades (**Fig. 3A**), generally corresponding to differences in genomic organization. One notable exception is Asclepias syriaca virus 2, which lacks additional ORFs yet clusters with betanucleorhabdoviruses that possess an accessory ORF between the M and G genes (**Fig. 3B**). On the other hand, within the major clade of viruses with the “basic” genomic organization, monocot-infecting viruses do not consistently form a monophyletic group. For instance, the two viruses associated to *Eichhornia* and *Stratiotes* group together, but the virus associated with the fern *Osmolindsaea* does not - instead, it groups with a virus infecting a dicot (**Fig. 3A**). This pattern suggests a lack of clear host-virus co-evolutionary trajectories for monocot- and fern-infecting betanucleorhabdoviruses, an observation consistent with previous findings in both invertebrate and vertebrate rhabdoviruses (Geoghegan et al., 2017), as well as in alphacytorhabdoviruses (Bejerman et al., 2023).

Based on current species demarcation criteria, viruses assigned to different species within the genus have a nt sequence identity lower than 75% in the complete genome sequence and occupy different ecological niches as evidenced by differences in host species and/or arthropod vectors (Walker et al., 2022). Therefore, the 25 novel betanucleorhabdoviruses identified in this study should be classified as new species in the genus *Betanucleorhabdovirus*.

### 3.4 Gammanucleorhabdovirus

The full-length coding regions of four novel gammanucleorhabdovirus genomes were assembled in this study (**Table 1**), increasing the number of putative members of this genus by 2-fold. These newly identified viruses were associated with four plant host species, all of which are woody dicots (**Table 3**). In contrast, the two gammanucleorhabdoviruses reported to date infect gramineous monocots (Redinbaugh et al., 2002; Alvarez-Quinto et al., 2022). This striking host difference suggests that the newly discovered viruses have undergone a host adaptation process favoring woody dicots during their evolutionary history. Thus, it is tempting to speculate that monocot-associated gammanucleorhabdoviruses have one type of vector, which is a leafhopper (Dietzgen et al., 2020); while those woody dicot-associated gammanucleorhabdoviruses have a distinct type of vector; that needs to be identified.

The genomic organization of the four novel gammanucleorhabdoviruses is 3′-N-P–P3-M-G-L-5′ (**Table 3, Fig. 4b**), which differs from that of the previously reported monocot-associated gammanucleorhabdoviruses, which contain an additional ORF between the third and fifth ORFs (Redinbaugh et al., 2002; Alvarez-Quinto et al., 2022) (**Fig. 4B**).

**Figure 4.**
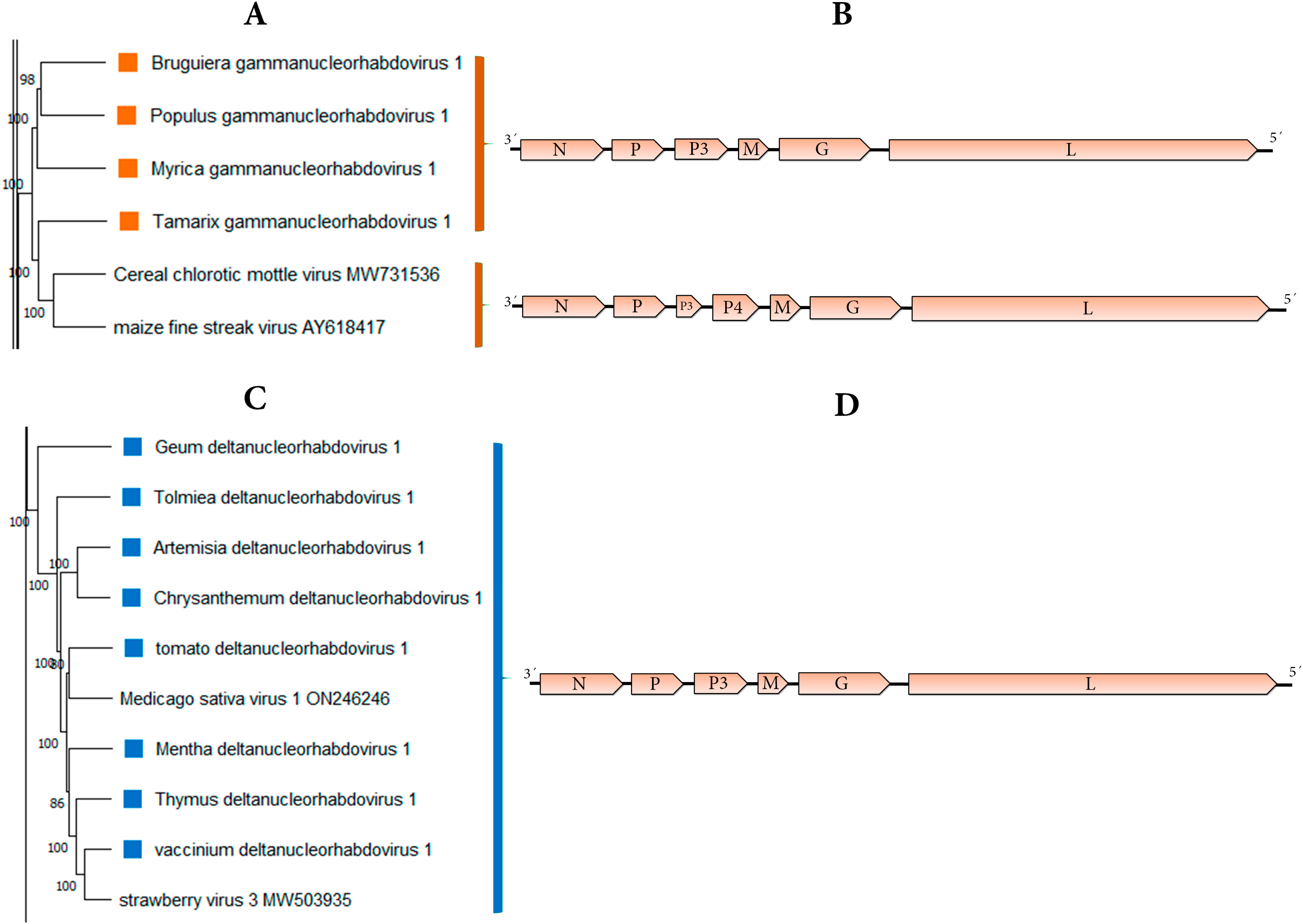
A: The maximum-likelihood phylogenetic tree shown in Fig. 1 was cropped to show only those viruses included in the genera *Gammanucleorhabdovirus* and *Deltanucleorhabdovirus*. The gammanucleorhabdoviruses and deltanucleorhabdoviruses identified in this study are noted with orange and blue rectangles, respectively. B: genomic organization of the viral sequences used in the phylogeny.

Despite differences in host type and genomic structure, the consensus gene junction sequences of the novel gammanucleorhabdoviruses are highly conserved among themselves and closely resemble those of the previously described members of the genus (**Table 5**).

Pairwise nt sequence identity values between the complete-coding genomes of the four novel viruses and the known gammanucleorhabdoviruses ranged between 58.7% and 61.9% (**Table S4**), with the highest values remaining below 62%. This relatively low sequence similarity suggests the possible existence of unexplored viral diversity, or “dark matter”, within the *Gammanucleorhabdovirus* genus.

Phylogenetic analysis based on the L protein aa sequence showed that the four novel viruses grouped with the two known gammanucleorhabdoviruses in a distinctive clade (**Fig. 1**). Within this clade, one subgroup includes the two monocot-associated viruses-cereal chlorotic mottle virus and maize fine streak virus-while three of the woody dicot-associated viruses formed a separate group, and the fourth woody dicot-associated virus formed its own monophyletic branch (**Fig. 4A**). Thus, gammanucleorhabdoviruses have a complex evolutionary history with distinct evolutionary trajectories according to the associated hosts, suggesting a shared host-virus evolution. Moreover, it is tempting to speculate that the ancestor gammanucleorhabdovirus was adapted to infect woody plants, while cereal chlorotic mottle virus and maize fine streak virus acquired an additional ORF during its adaptation to their monocot host.

Consistent with established species demarcation criteria, viruses assigned to different species within the genus have a nt sequence identity lower than 75% in the complete genome sequence and occupy different ecological niches as evidenced by differences in host species and/or arthropod vectors (Walker et al., 2022). Therefore, the four novel gammanucleorhabdoviruses identified in this study should be classified as new species in the genus *Gammanucleorhabdovirus*.

### 3.5 Deltanucleorhabdovirus

The full-length coding regions of eight novel deltanucleorhabdoviruses were assembled in this study (**Table 4**), increasing the number of putative members of this genus by 4-fold. The viruses detected were associated with eight plant host species (**Table 4**). All the host plants were herbaceous dicots, like the two deltanucleorhabdoviruses reported to date (Li et al., 2022B; Medberry et al., 2022). Therefore, this group of viruses likely had a host adaptation trajectory to infect herbaceous dicots during their evolution that may be likely linked to the host preference of their insect vector, as this is a strong correlation among plant rhabdoviruses (Dietzgen et al., 2020).

The eight novel deltanucleorhabdovirus genomes identified in this study exhibit a conserved genomic organization of 3′-N-P–P3-M-G-L-5′ (**Table 4, Fig. 4b**), consistent with the two previously characterized deltanucleorhabdoviruses (Li et al., 2022B; Medberry et al., 2022). No accessory ORFs were detected in any of these genomes. This conserved genomic structure suggests that deltanucleorhabdoviruses likely share a recent common ancestor and have not acquired additional genes during their evolution. This lineage may represent a relatively recent divergence from the ancestral nucleorhabdovirus clade. Furthermore, the consensus gene junction sequences of the novel deltanucleorhabdoviruses are highly similar, both among themselves and with those of previously reported deltanucleorhabdoviruses (**Table 5**), further supporting a shared evolutionary origin.

Pairwise nt sequence identity values between the complete-coding genome of the eight novel viruses and the known deltanucleorhabdoviruses range between 58.3% and 70.7% (**Table S5**). This level of divergence suggests that the genus is relatively compact, with limited uncharacterized diversity and not a large reservoir of “viral dark matter.”

The phylogenetic analysis based on the L protein aa sequences places the eight novel viruses together with the two known deltanucleorhabdoviruses in a distinctive clade (**Fig. 1**). Within this group of ten viruses, four subclades are evident, but all are separated by relatively short branches (**Fig. 4A**), consistent with a relatively recent diversification.

Viruses assigned to different species within the genus have a nucleotide sequence identity lower than 75% in the complete genome sequence and occupy different ecological niches as evidenced by differences in host species and/or arthropod vectors (Walker et al., 2022). Therefore, the eight novel deltanucleorhabdoviruses identified in this study should be classified as new species in the genus *Deltanucleorhabdovirus*.

### 3.6. Strengths and Limitations of sequence discovery through data mining

As previously demonstrated by Bejerman and colleagues (Bejerman et al., 2022; 2023), independent validation through re-analyzing of SRA data can significantly enhance our understanding of RNA virus genomes with unique features. However, a key limitation of this approach is the lack of access to original biological material, which impedes replication and verification of the assembled viral genome sequences, an inherent weakness of data mining-based virus discovery. Additionally, potential issues such as contamination, low sequencing quality, sample spill-over, and other technical artifacts present a risk of generating false-positive, chimeric assemblies, or incorrect host assignments. Consequently, caution must be exercised when interpreting results derived from publicly available SRA data. To strengthen and validate our findings, we strongly recommend generating new RNAseq datasets from the predicted plant hosts. While our approach has inherent limitations, some aspects of our virus discovery strategy, such as iterative contig extension, virus detection in independent datasets, and comprehensive sequence validation, can aid to alleviate some of these challenges and provide further evidence for identification. Nevertheless, it is key to emphasize that all virus-host associations and detections should be considered preliminary until confirmed through further investigation.

## 4. Conclusions

This study is the final chapter in our trilogy exploring the cryptic diversity of plant-associated rhabdoviruses through data mining, highlighting the value of analyzing publicly available SRA data., This strategy has proven to be a powerful tool not only for accelerating the discovery of novel rhabdoviruses but also for deepening our understanding of their evolutionary history and refining their taxonomic classifications. Employing this approach, we uncovered a hidden diversity of nucleorhabdovirus-like sequences, effectively doubling the number of known members in this group. Our findings also expanded their known host range, including the first nucleorhabdovirus associated with ferns and the first gammanucleorhabdoviruses associated with dicotyledonous hosts. These discoveries shed light on the complexity of their evolutionary dynamics despite the limited variation observed in their genomic organization. Future studies should assess diverse aspects of the biology and ecology of the novel viruses identified in this study, including their intracellular localization, their potential to induce symptoms in their hosts, and their likely arthropod vectors.

## Acknowledgments

We would like to express genuine appreciation to the producers of the original sequence data used for this study, which are cited in **Tables 1, 2, 3 and 4**. By ensuing open science practices with accessible raw sequence data in open public repositories, these authors supported contributions based on secondary data analyses.

## Author Contributions

Conceptualization, N.B and H.D; data analysis, N.B and H.D; writing—original draft preparation, N.B; writing—review and editing, N.B, R.G.D and H.D. All authors have read and agreed to the published version of the manuscript.

## Institutional Review Board Statement

Not applicable for studies not involving humans or animals.

## Informed Consent Statement

Not applicable for studies not involving humans.

## Data availability statement

Nucleotide sequence data reported are available in the Third Party Annotation Section of the DDBJ/ENA/GenBank databases under the accession numbers TPA: BK070473-BK070520 and can be found as **Supplementary Material 1** of this submission.

