## Supplementary material for "Expanding the known nucleorhabdovirus world: the final chapter in a trilogy exploring the hidden diversity of plant-associated rhabdoviruses": Table 1

**Table 1**. Summary of novel alphanucleorhabdoviruses identified from plant RNA-seq data available on NCBI.

| **Plant host** | **Taxa/**  **family** | **Virus name/**  **Abbreviation** | **Bioproject ID/**  **Data citation** | **Length (nt)** | **Accession number** | **Protein ID** | **Length (aa)** | **Highest scoring virus- protein/*E*-value/query coverage%/identity% (Blast P)** |
| --- | --- | --- | --- | --- | --- | --- | --- | --- |
| black chokeberry  (*Aronia melanocarpa*) | Dicot/  *Rosacaeae* | Aronia  alphanucleorhabdovirus 1/  AroANRV1 | PRJNA603127/  Mahoney et al., (2022) | 13487 | BK070473 | N  P  P3  M  G  L | 479  293  279  243  590  1936 | PeV1-N/0.0/99/59.03  PeV1-P/8e-68/99/41.22  PeV1-P3/6e-106/99/53.79  PeV1-M/73-61/96/45.11  PeV1-G/0.0/98/49.06  PeV1-L/0.0/99/60.94 |
| Atacama azorella  (*Azorella atacamensis*) | Dicot/  *Apiaceae* | Azorella  alphanucleorhabdovirus 1/  AzoANRV1 | PRJNA687835/  Eshel et al., (2021) | 13516 | BK070474 | N  P  P3  M  G  L | 457  316  287  237  597  1947 | ATV1-N/5e-152/89/51.58  ATV1-P/6e-15/76/24.16  ATV1-P3/6e-13/68/23.53  ATV1-M/7e-05/63/21.52  ATV1-G/1e-121/92/35.23  ATV1-L/0.0/98/46.77 |
| caper bush  (*Capparis spinosa*) | Dicot/  *Capparaceae* | Capparis  alphanucleorhabdovirus 1/  CapANRV1 | PRJNA792936/  Wang et al., (2022) | 13727 | BK070475 | N  P  P3  M  G  L | 495  321  306  239  592  1937 | RYSV-N/6e-57/87/30.65  No hits  No hits  EMDV-M/1e-10/83/22.49  PYDV-G/9e-61/90/29.27  XARV-L/0.0/77/44.11 |
| Cassava  (*Manihot esculenta*) | Dicot/  *Euphorbiaceae* | Cassava  alphanucleorhabdovirus 1/  CasANRV1 | PRJNA578024/  Hu et al., (2021) | 13729 | BK070476 | N  P  P3  M  G  L | 492  326  284  251  614  1951 | JYBaV-N/2e-102/93/37.93  XARV-P/3e-05/57/28.06  PYDV-P3/3e-48/97/31.77  PhMCoV-M/5e-17/82/28.5  PhCMoV-G/2e-122/87/35.36  BabRV1-L/0.0/99/44.76 |
| bermudagrass  (*Cynodon dactylon*) | Monocot/  *Poaceae* | Cynodon  alphanucleorhabdovirus 1/  CynANRV1 | PRJNA693979/  Li et al., (2022A) | 12666 | BK070477 | N  P  P3  M  G  L | 443  272  289  233  595  1926 | MIMV-N/0.0/100/71.91  MIMV-P/7e-122/100/63.37  MIMV-P3/5e-114/96/52.86  MIMV-M/2e-97/100/60.09  MIMV-G/0.0/96/60.63  MIMV-L/0.0/98/74.71 |
| bedgrass  (*Hildaea pallens*) | Monocot/  *Poaceae* | Hildaea  alphanucleorhabdovirus 1/  HilANRV1 | PRJNA756291/  Huang et al., (2022) | 14636 | BK070478 | N  P  P3  M  G  P6  L | 476  326  295  275  656  130  1962 | RYSV-N/0.0/86/68.36  RYSV-P/7e-116/99/55.52  RYSV-P3/2e-159/98/69.18  RYSV-M/4e-115/99/61.17  WYSV-G/0.0/90/62.25  No hits  WYSV-L/0.099/67.99 |
| ryegrass  (*Lolium perenne*) | Monocot/  *Poaceae* | Lolium alphanucleorhabdovirus 1/  LolANRV1 | PRJEB30432/  Maus et al., (2020) | 14831 | BK070479 | N  P  P3  M  G  P6  L | 480  331  289  272  661  104  1963 | WYSV-N/0.0/85/62.59  RYSV-P/2e-70/96/39.44  RYSV-P3/1e-79/97/42.96  RYSV-M/1e-92/100/50.37  WYSV-G/0.0/98/52.27  No hits  WYSV-L/0.0/100/64.53 |
| water bluebell  (*Ruellia longepetiolata*) | Dicot/  *Acanthaceae* | Ruellia  alphanucleorhabdovirus 1/  RueANRV1 | PRJNA323650/  Zhuang et al., (2017) | 13531 | BK070480 | N  X  P  P3  M  G  L | 487  115  329  289  255  615  1945 | PhCMoV-N/5e-91/96/36  No hits  XARV-P/3e-04/39/28.89  XARV/8e-31/85/27.02  EMDV-M/2e-16/78/26.24  EMDV-G/2e-110/94/33.17  PhCMoV-L0.0/99/45.03 |
| Sprawling schiedea  (*Schiedea hookeri*) | Dicot/  *Caryophyllaceae* | Schiedea  alphanucleorhabdovirus 1/  SchANRV1 | PRJNA491458/  Nevado et al., (2019) | 12872 | BK070481 | N  P  P3  M  G  L | 499  317  321  254  611  1887 | RYSV-N/5e-55/88/30.36  No hits  EMDV-P3/1e-05/71/22.27  PhCMoV-M/7e-12/88/26.02  PhCMoV-G/1e-56/88/28.83  EMDV-L/0.0/98/38.03 |
| fireweed  (*Senecio magnificus*) | Dicot/  *Asteraceae* | Senecio alphanucleorhabdovirus 1/  SenANRV1 | PRJNA371565/  Jayasena et al., (2017) | 13130 | BK070482 | N  P  P3  M  G  L | 458  312  314  231  610  1929 | ATV1-N/1e-99/89/39.85  No hits  No hits  No hits  ATV1-G/2e-100/84/33.4  ATV1-L/0.0/98/37.09 |
