## Supplementary material for "Expanding the known nucleorhabdovirus world: the final chapter in a trilogy exploring the hidden diversity of plant-associated rhabdoviruses": Table 2

**Table 4**. Summary of novel betanucleorhabdoviruses identified from plant RNA-seq data available on NCBI.

| **Plant host** | **Taxa/**  **family** | **Virus name/**  **Abbreviation** | **Bioproject ID/**  **Data citation** | **Length (nt)** | **Accession number** | **Protein ID** | **Length (aa)** | **Highest scoring virus- protein/*E*-value/query coverage%/identity% (Blast P)** |
| --- | --- | --- | --- | --- | --- | --- | --- | --- |
| Sagebrush  (*Artemisia sieversiana*) | Dicot/  *Asteraceae* | Artemisia  betanucleorhabdovirus 1/  ArtBNRV1 | PRJNA834888/  Zhang et al., (2022) | 14459 | BK070483 | N  P  P3  M  G  L | 471  365  327  280  664  2149 | PBNRV1-N/7e-155/94/47.48  PBNRV1-P/2e-25/97/25.99  PBNRV1-P3/1e-84/99/40.18  CnV1-M/3e-27/94/30.22  PBNRV1-G/0.0/97/53.26  PBNRV1-L/0.0/99/54.78 |
| Ramie  (*Boehmeria nivea*) | Dicot/  *Urticaceae* | Boehmeria  betanucleorhabdovirus 1/  BoeBNRV1 | PRJNA252411/  Chen et al., (2014) | 13766 | BK070484 | N  P  P3  M  G  L | 458  345  324  283  632  2107 | BNRV1-N/0.0/99/69.45  CIRV-P/2e-88/93/46.48  ToBNRV2-P3/3e-163/99/64.09  BNRV1-M/9e-94/94/52.21  ToBNRV2-G/0.0/99/60.7  SaBNV4-L/0.0/99/58.53 |
| golden camellia  (*Camellia nitidissima*) | Dicot/  *Theaceae* | Camellia  betanucleorhabdovirus 1/  CamBNRV1 | PRJNA794001/  Liu et al., (2022) | 14052 | BK070485 | N  P  P3  M  U  G  L | 444  356  315  289  92  623  2052 | AaNV-N/0.0/99/57.34  AaNV-P/7e-29/96/30.17  AaNV-P3/3e-48/32.44/93  AaNV-M/3e-13/68/27.86  No hits  AaNV-G/3e-180/95/45.58  AaNV-L/0.0/99/48.79 |
| canola  (*Brassica napus*) | Dicot/  *Brassicaceae* | canola betanucleorhabdovirus 1/  CanBNRV1 | PRJNA881481/  Mahdikhani, R., WA University, Australia, unpublished | 14019 | BK070486 | N  P  P3  M  G  L | 468  349  327  284  647  2131 | SaBNV2-N/2e-180/97/56.52  TarBNRV1-P/1e-31/92/27.52  CnV1-P3/6e-69/98/35.69  BCaRV-M/5e-50/79/40.27  CnV1-G/0.0/88/45.88  CnV1-L/0.0/99/47.3 |
| Korean bittercress  (Cardamine leucantha) | Dicot/  *Brassicaceae* | Cardamine betanucleorhabdovirus 1/  CarBNRV1 | PRJDB9421/  Araki et al., (2020) | 13800 | BK070487 | N  P  P3  M  G  L | 458  345  324  282  635  2109 | SaBNV2-N/0.0/99/67.61  ToBNRV2-P/1e-77/97/41.37  ToBNRV2-P3/1e-165/99/64.71  ToBNRV2-M/3e-95/96/51.84  SaBNV2-G/0.0/98/59.84  SaBNV2/0.0/99/58.2 |
| thick-leaved mum  (*Chrysanthemum crassum*) | Dicot/  *Asteraceae* | Chrysanthemum betanucleorhabdovirus 1/  ChrBNRV1 | PRJNA380725/  Guan et al., (2017) | 14675 | BK070488 | N  P  P3  M  G  L | 468  334  326  285  660  2141 | PBNRV1-N/1e-161/99/46.76  PBNRV1-P/1e-31/98/27.42  PBNRV1-P3/1e-73/94/38.96  BCaRV-M/1e-34/81/37.5  PBNRV1-G/0.0/98/53.92  PBNRV1-L/0.0/99/54.46 |
| golden thread root  (*Coptis teeta*) | Dicot/  *Ranunculaceae* | Coptis betanucleorhabdovirus 1/  CopBNRV1 | PRJNA433257/  He et al., (2018) | 13486 | BK070489 | N  P  P3  M  G  L | 463  322  322  271  642  2108 | GSPNuV-N/0.0/96/64.96  GSPNuV-P/1e-69/95/38.62  BFTaV-P3/4e-120/99/47.96  GSPNuV-M/2e-86/91/51.98  GSPNuV-G/0.0/90/62.63  GSPNuV/0.0/99/54.96 |
| Fried-egg orchid  (*Dendrobium chrysotoxum*) | Monocot/  *Orchidaceae* | Dendrobium  betanucleorhabdovirus 1/  DenBNRV1 | PRJNA691441/  Zhang et al., (2021) | 13826 | BK070490 | N  P  P3  M  G  L | 461  355  321  280  636  2101 | SYVV-N/0.0/100/59.79  GSPNuV-P2e-69/96/37.54  SYVV-P3/2e-100/100/44.24  SYVV-M/5e-66/97/43.57  SYVV-G/0.0/99/51.96  SYVV-L/0.0/100/54.15 |
| Dicot plant | - | dicot associated  betanucleorhabdovirus 1/  DaBNRV1 | PRJNA375958/  Wu et al., (2018) | 13542 | BK070491 | N  P  P3  M  G  L | 463  366  321  279  659  2097 | SYVV-N/0.0/99/60.13  GSPNuV-P/2e-58/97/33.06  GSPNuV-P3/2e-85/98/39.5  SYVV-M/1e-63/93/39.85  SYVV-G/0.0/95/50.71  SYVV-L/0.0/100/53.17 |
| Brazilian water hyacinth  (*Eichhornia paniculata*) | Monocot/  *Pontedieraceae* | Eichhornia  betanucleorhabdovirus 1/  EicBNRV1 | PRJNA266681/  University of Toronto, Canada, unpublished | 13818 | BK070492 | N  P  P3  M  G  L | 472  318  318  275  626  2099 | GSPNuV-N/3e-174/96/55.04  GSPNuV-P/7e-47/95/31.14  GSPNuV-P3/3e-82/94/41.25  BFTaV-M/4e-63/86/42.02  BmV2-G/0.0/96/46.17  BFTaV-L/0.0/99/47.92 |
| swamp candles  (*Lysimachia terrestris*) | Dicot/  *Primulaceae* | Lysimachia  betanucleorhabdovirus 1/  LysBNRV1 | PRJNA422719/  Marx et al., (2021) | 16292 | BK070493 | N  P  P3  M  U  G  L | 465  347  307  311  242  640  2115 | ApRVA-N/0.0/98/60.09  ApRVA-P/7e-30/96/26.55  ApRVA-P3/2e-90/95/47.3  ApRVA-M/6e-42/85/32.22  No hits  ApRVA-G/0.0/92/50.99  ApRVA-L/0.0/99/51.18 |
| Canada mint  (*Mentha canadensis*) | Dicot/  *Lamiaceae* | Mentha  betanucleorhabdovirus 1/  MenBNRV1 | PRJNA724910/  Yu et al., (2021) | 13455 | BK070494 | N  P  P3  M  G  L | 468  324  330  308  643  2108 | CnV1-N/0.0/99/54.72  CnV1-P/83-62/98/35.71  CnV1-P3/4e-92/94/43.27  CnV1-M/5e-5/89/36  CnV1-G/0.0/98/52.26  CnV1-L/0.0/99/52.65 |
| Moyeam  (*Nekemias grossedentata*) | Dicot/  *Vitaceae* | Nekemias  betanucleorhabdovirus 1/  NekBNRV1 | PRJNA777899/  Zhang, J., Fudan University, China, unpublished | 14151 | BK070495 | N  P  P3  M  U  G  L | 443  350  315  295  95  628  2049 | AaNV-N/0.0/99/56.33  AaNV-P/8e-24/90/27.03  AaNV-P3/6e-41/93/29.05  AaNV-M/3e-09/67/23.74  No hits  AaNV-G/9e-177/98/43.88  AaNV-L/0.0/99/49 |
| Proliferous water lily  (*Nymphaea prolifera*) | Dicot/  *Nymphaeaceae* | Nymphaea betanucleorhabdovirus 1/  NymBNRV1 | PRJNA565347/  Zhang et al., (2020) | 13519 | BK070496 | N  P  P3  M  G  L | 472  327  318  279  629  2098 | PleArV1-N/2e-179/93/58.41  BCaRV-P/3e-45/97/32.09  GSPNuV-P3/4e-81/96/37.99  GSPNuV-M/2e-63/94/40.15  BmV2-G/0.0/96/47.49  BFTaV-L/0.0/99/49.48 |
| Screw fern  (*Osmolindsaea odorata*) | *Polypodiophyta* /  *Lindsaeaceae* | Osmolindsaea  betanucleorhabdovirus 1/  OsmBNRV1 | PRJNA281136/  Shen et al., (2018) | 13265 | BK070497 | N  P  P3  M  G  L | 450  311  322  272  632  2094 | GSPNuV-N/0.0/97/61.68  PleArV1-P/9e-72/97/40.12  GSPNuV-P3/4e-113/100/47.83  GSPNuV-M/5e-81/86/50  GSPNuV-G/0.0/94/53.6  BFTaV-L/0.0/99/52.09 |
| dwarf snow rhododendron  (*Rhododendron nivale*) | Dicot/  *Ericaceae* | Rhododendron  betanucleorhabdovirus 2/  RhoBNRV2 | PRJNA540086/  Hao, D., China. Unpublished | 14196 | BK070498 | N  P  P3  M  G  L | 464  352  326  286  647  2116 | RhoDeV1-N/0-.0/100/62.58  RhoDeV1-P/4e-112/98/49.43  RhoDeV1-P3/2e-139/99/56.79  RhoDeV1-M/4e-92/94/49.64  RhoDeV1-G/0.0/97/63.17  RhoDeV1-L/0.0/94/67.05 |
| Turkish ricotia  (*Ricotia aucheri*) | Dicot/  *Brassicaceae* | Ricotia  betanucleorhabdovirus 1/  RicBNRV1 | PRJNA634714/  Guo et al., (2020) | 13688 | BK070499 | N  P  P3  M  G  L | 460  349  323  283  620  2103 | SYVV-N/0.0/98/58.84  GSPNuV-P/2e-53/97/34.19  GSPNuV-P3/1e-96/98/43.26  SYVV-M/9e-74/91/43.68  BFTaV-G/0.0/95/51.17  BFTaV-L/0.0/99/53.89 |
| Creeping yellowcress fieldcress  (*Rorippa sylvestri*s) | Dicot/  *Brassicaceae* | Rorippa betanucleorhabdovirus 1/  RorBNRV1 | PRJEB73992/  Van Veen et al., (2024) | 13439 | BK070500 | N  P  P3  M  G  L | 459  365  327  281  635  2109 | PleArV1-N/0.0/96/61.09  PleArV1-P/1e-57/95/31.99  GSPNuV-P3/2e-92/91/42  BFTaV-M/73-70/84/44.21  GSPNuV-G/0.0/96/55.75  BFTaV-L/0.0/99/52.74 |
| purple willow  (*Salix purpurea*) | Dicot/  *Salicaceae* | Salix  betanucleorhabdovirus 1/  SalBNRV1 | PRJEB18638/  Yanitch et al., (2017) | 14195 | BK070501 | N  P  P3  M  G  L | 463  365  330  271  663  2103 | SaBNV4-N/2e-130/99/44.44  PBNRV1-P/2e-25/92/25.9  RhoDeV1-P3/3e-44/95/26.69  CnV1-M/1e-20/79/29.49  BmV2-G/9e-114/85/34.95  BFTaV-L/0.0/99/43.94 |
| red sage  (*Salvia miltiorrhiza*) | Dicot/  *Lamiaceae* | Salvia betanucleorhabdovirus 1/  SalvBNRV1 | PRJNA795600/  Cui et al., (2021) | 13722 | BK070502 | N  P  P3  M  G  L | 460  344  322  298  646  2101 | BFTaV-N/0.0/99/69.43  BFTaV-P/5e-97/99/45.45  BFTaV-P3/2e-141/99/57.81  BFTaV-M/8e-103/85/55.47  BFTaV-G/0.0/97/68.04  BFTaV-L/0.0/100/60.04 |
| Water soldier  (*Stratiotes aloides*) | Monocot/  *Hydrocharitaceae* | Stratiotes  betanucleorhabdovirus 1/  StrBNRV1 | PRJNA809041/  Chen et al., (2022A) | 13637 | BK070503 | N  P  P3  M  G  L | 472  320  318  276  639  2103 | PleArV1-N/6e-177/88/58.03  GSPNuV-P/7e-45/98/31.21  GSPNuV-P3/3e-82/94/39.4  BFTaV-M/6e-65/92/38.91  SaBNV2-G/0.0/96/44.86  SaBNV3-L/0.099/49.69 |
| five-ribbed thyme  (*Thymus quinquecostatu*s) | Dicot/  *Lamiaceae* | Thymus betanucleorhabdovirus 1/  ThyBNRV1 | PRJNA690675/  Sun et al., (2022) | 13779 | BK070504 | N  P  P3  M  G  L | 460  347  322  297  646  2104 | BFTaV-N/0.0/97/70  BFTaV-P/5e-103/100/47.18  BFTaV-P3/1e-151/99/60.5  BFTaV-M/3e-110/95/55.14  BFTaV-G/0.0/97/66.3  BFTaV-L/0.0/100/59.54 |
| truncate gourd  (*Trichosanthes truncata*) | Dicot/  *Cucurbitaceae* | Trichosanthes  betanucleorhabdovirus 1/  TriBNRV1 | PRJNA624798/  Guo et al., (2020) | 13952 | BK070505 | N  P  P3  M  G  L | 458  356  324  278  631  2108 | BNRV1-N/0.0/99/70.77  ToBNRV2-P/5e-96/94/48.52  SaBNV2-P3/9e-157/99/62.93  BNRV1-M/2e-105/88/61.13  ToBNRV2-G/0.0/99/58.37  SaBNV4-L/0.0/100/57.73 |
| Violet  (*Viola baoshanensis*) | Dicot/  *Violaceae* | Viola  betanucleorhabdovirus 1/  VioBNRV1 | PRJNA524759/  Shu et al., (2019) | 13469 | BK070506 | N  P  P3  M  G  L | 468  323  320  278  633  2109 | PyRV1-N/0.0/99/76.01  PyRV1-P/8e-141/98/62.62  PyRV1-P3/7e-167/99/65.41  PyRV1-M/5e-110/91/58.98  PyRV1-G/0.0/99/73.45  PyRV1-L/0.0/99/67.57 |
| Wild basil  (*Ziziphora clinopodioides*) | Dicot/  *Lamiaceae* | Ziziphora betanucleorhabdovirus 1/  ZizBNRV1 | PRJNA566012/  He et al., (2020) | 13778 | BK070507 | N  P  P3  M  G L | 460  345  345  301  634  2104 | BFTaV-N/0.0/97/69.33  BFTaV-P/4e-100/100/46.39  BFTaV-P3/8e-152/99/60  BFTaV-M/2e-104/99/50.33  BFTaV-G/0.0/98/68.15  BFTaV-L/0.0/100/60.31 |
