## Supplementary material for "Expanding the known nucleorhabdovirus world: the final chapter in a trilogy exploring the hidden diversity of plant-associated rhabdoviruses": Table 3

**Table 3**. Summary of novel gammanucleorhabdoviruses identified from plant RNA-seq data available on NCBI.

| **Plant host** | **Taxa/**  **family** | **Virus name/**  **Abbreviation** | **Bioproject ID/**  **Data citation** | **Length (nt)** | **Accession number** | **Protein ID** | **Length (aa)** | **Highest scoring virus- protein/*E*-value/query coverage%/identity% (Blast P)** |
| --- | --- | --- | --- | --- | --- | --- | --- | --- |
| oriental mangrove  (*Bruguiera gymnorhiza*) | Dicot/  *Rhizophoraceae* | Bruguiera  gammanucleorhabdovirus 1/  BruGNRV1 | PRJNA817364/  He et al., (2022) | 13259 | BK070517 | N  P  P3  M  G  L | 468  355  323  228  591  1947 | MFSV-N/1e-90/82/38.73  MFSV-P/3e-10/81/24.51  No hits  MFSV-M/2e-10/76/26.29  MFSV-G/3e-106/91/34.29  MFSV-L/0.0/98/42.4 |
| Chinese bayberry  (*Myrica rubra*) | Dicot/  *Myricaceae* | Myrica  gammanucleorhabdovirus 1/  MyrGNRV1 | PRJNA368720/  Shi et al., (2018) | 14282 | BK070518 | N  P  P3  M  G  L | 469  378  312  248  600  1966 | CCMoV-N/5e-90/90/37.47  No hits  CCMoV-P3/8e-11/88/22.96  MFSV-M/4e-11/74/26.34  MFSV-G/1e-92/92/32.04  MFSV-L/0.0/96/41.83 |
| black Cottonwood  (*Populus trichocarpa*) | Dicot/  *Salicaceae* | Populus gammanucleorhabdovirus 1/  PopGNRV1 | PRJNA568075/  Lenz et al., (2021) | 14326 | BK070519 | N  P  P3  M  G  L | 469  387  324  236  591  1941 | MFSV-N/9e-100/97/37.61  MFSV-P/0.012/21/27.06  No hits  No hits  MFSV-G/4e-113/91/35.53  MFSV-L/0.0/99/42.15 |
| Tamarisk  (*Tamarix ramosissima*) | Dicot/  *Tamaricaceae* | Tamarix gammanucleorhabdovirus 1/  TamGNRV1 | PRJNA799438/  Chen et al., (2022B) | 13791 | BK070520 | N  P  P3  M  G  L | 481  354  322  222  604  1952 | MFSV-N/3e-108/85/40.1  MFSV-P/1e-08/76/24.58  No hits  CCMoV-M/4e-10/86/24.76  MFSV-G/1e-107/96/33.9  MFSV-L/0.0/100/46.18 |
