## Supplementary material for "Expanding the known nucleorhabdovirus world: the final chapter in a trilogy exploring the hidden diversity of plant-associated rhabdoviruses": Table 4

**Table 4**. Summary of novel deltanucleorhabdoviruses identified from plant RNA-seq data available on NCBI.

| **Plant host** | **Taxa/**  **family** | **Virus name/**  **Abbreviation** | **Bioproject ID/**  **Data citation** | **Length (nt)** | **Accession number** | **Protein ID** | **Length (aa)** | **Highest scoring virus- protein/*E*-value/query coverage%/identity% (Blast P)** |
| --- | --- | --- | --- | --- | --- | --- | --- | --- |
| sweet wormwood  (*Artemisia annua*) | Dicot/  *Asteraceae* | Artemisia  deltanucleorhabdovirus 1/  ArtDNRV1 | PRJNA752933/  Ma et al., (2021) | 15227 | BK070508 | N  P  P3  M  G  L | 519  293  321  268  690  1998 | MsV1-N/0.0/91/56.51  StrV3-P/6e-68/83/41.63  StrV3-P3/1e-101/100/46.11  StrV3-M/3e-25/86/29.18  StrV3-G/0.0/87/54.21  StrV3-L/0.0/99/58.48 |
| thick-leaved mum  (*Chrysanthemum crassum*) | Dicot/  *Asteraceae* | Chrysanthemum deltanucleorhabdovirus 1/  ChrDNRV1 | PRJNA380725/  Guan et al., (2017) | 14830 | BK070510 | N  P  P3  M  G  L | 520  291  321  257  691  1994 | StrV3-N/4e-180/91/54.09  StrV3-P/1e-57/88/39.38  StrV3-P3/4e-106/99/48.44  MsV1-M/4e-33/97/31.68  MsV1-G/0.0/97/51.41  StrV3-L/0.0/96/59.81 |
| Herb bennet  (*Geum urbanum*) | Dicot/  *Rosaceae* | Geum  deltanucleorhabdovirus 1/  GeuDNRV1 | PRJEB23354/  (Jordan et al., 2018) | 14140 | BK070511 | N  P  P3  M  G  L | 474  302  319  281  814  1985 | StrV3-N/1e-44/93/28.57  No hits  StrV3-P3/2e-27/87/24.91  No hits  StrV3-G/6e-44/80/24.19  MsV1-L/0.0/84/42.33 |
| spearmint  (*Mentha spicata*) | Dicot/  *Lamiaceae* | Mentha  deltanucleorhabdovirus 1/  MenDNRV1 | PRJDB2527/  Jin et al., (2014) | 14557 | BK070512 | N  P  P3  M  G  L | 512  298  322  255  700  1999 | StrV3-N/0.0/94/60.78  StrV3-P/1e-104/91/52.73  StrV3-P3/2e-171/98/69.5  StrV3-M/6e-57/86/47.27  StrV3-G/0.0/88/60.84  StrV3-L/0.0/99/66.18 |

| five-ribbed thyme  (*Thymus quinquecostatu*s) | Dicot/  *Lamiaceae* | Thymus deltanucleorhabdovirus 1/  ThyDNRV1 | PRJNA690675/  Sun et al., (2022) | 13988 | BK070513 | N  P  P3  M  G  L | 501  292  322  251  675  1986 | StrV3-N/0.0/96/70.45  StrV3-P/1e-118/90/59.62  StrV3-P3/0.0/99/79.06  StrV3-M/2e-81/89/60.53  StrV3-G/0.0/90/70.82  StrV3-L/0.0/99/72.85 |
| --- | --- | --- | --- | --- | --- | --- | --- | --- |
| Piggyback plant  (*Tolmiea menziesii*) | Dicot/  *Saxifragaceae* | Tolmiea deltanucleorhabdovirus 1/  TolDNRV1 | PRJNA507776/  Visger et al., (2019) | 14165 | BK070514 | N  P  P3  M  G  L | 529  290  324  249  692  1996 | StrV3-N/0.0/85/58.46  StrV3-P/3-e40/95/31.43  StrV3-P3/9e-112/99/49.85  MsV1-M/8e-34/97/31.01  StrV3-G/0.0/97/47.28  MsV1-L/0.0/94/56.69 |
| tomato  (*Solanum lycopersicum*) | Dicot/  *Solanaceae* | tomato deltanucleorhabdovirus 1/  TomDNRV1 | PRJNA526255/  Powell et al., (2022) | 14175 | BK070515 | N  P  P3  M  G  L | 495  289  329  245  687  1989 | MsV1-N/0.0/90/66.29  StrV3-P/3e-64/92/40.82  MsV1-P3/1e-158/96/63.84  MsV1-M/3e-59/98/40.86  MsV1-G/0.0/93/59.53  MsV1-L/0.0/96/65.46 |
| American cranberry  (*Vaccinium macrocarpon*) | Dicot/  *Ericaceae* | Vaccinium deltanucleorhabdovirus 1/  VacDNRV1 | PRJNA260125/  Sun et al., (2015) | 14415 | BK070516 | N  P  P3  M  G  L | 500  294  322  255  684  1987 | StrV3-N/0.0/96/82.1  StrV3-P/2e-158/92/72.53  StrV3-P3/0.0/100/87.58  StrV3-M/3e-103/91/69.1  StrV3-G/0.0/99/73.79  StrV3-L/0.0/99/79.51 |
