## Supplementary material for "Expanding the known nucleorhabdovirus world: the final chapter in a trilogy exploring the hidden diversity of plant-associated rhabdoviruses": Table 5

**Table 5**. Consensus conserved nucleorhabdovirus gene junction sequences

| **Genus** | **Virus*** | **3´end mRNA** | **intergenic spacer** | **5´end mRNA** |
| --- | --- | --- | --- | --- |
| *Alphanucleorhabdovirus* | AroANRV1 | AUUU(A/C)UUUU | G(N)n | UUG |
|  | AzoANRV1 | UUAAUUUUU | GGG | UUG |
|  | CapANRV1 | AUUCUUUUU | GGG | UUG |
|  | CasANRV1 | AUUAUUUUU | GGG | UUG |
|  | CynANRV1 | AUUCUUUUU | GGG | UUG |
|  | HilANRV1 | UUUAUUUUU | GGG | UUG |
|  | LolANRV1 | AUUAUUUUU | GGGG | UUG |
|  | RueANRV1 | AUUAUUUUU | GGG | UUG |
|  | SchANRV1 | AUUCUUUUU | GGG | UUG |
|  | SenANRV1 | AUUCUUUUU | GGG | UUG |
|  | ATV1 | AUUCUUUUU | GGG | UUG |
|  | ArtCaNV1 | AUUAUUUUU | GGG | UUG |
|  | BabRV1 | AUUAUUUUU | GGG | UUG |
|  | CYDV | AUUAUUUUU | GGG | UUG |
|  | EMDV | AUUAUUUUU | GGG | UUG |
|  | JYBaV | AUUAUUUUU | GGG | UUG |
|  | MIMV | AUUCUUUUU | GGG | UUG |
|  | MMV | AUUCUUUUU | GGG | UUG |
|  | MMaV | AUUCUUUUU | GGG | UUG |
|  | PhCMoV | AUUAUUUUU | GGG | UUG |
|  | PeV1 | AUUU(A/C)UUUU | G(N)n | UUG |
|  | PYDV | AUUAUUUUU | GGG | UUG |
|  | RYSV | AUUAUUUUU | GGG | UUG |
|  | TaVCV | AUUCUUUUU | GGG | UUG |
|  | TARV1 | AUUAUUUUU | GGG | UUG |
|  | WYSV | UAAAUUUUU | GGGG | UUG |
|  | XARV | AUUAUUUUU | GGG | UUG |
| *Betanucleorhabdovirus*  *Betanucleorhabdovirus* | ArtBNRV1 | AUUCUUUUU | GG | UUG |
|  | BoeBNRV1 | AUUCUUUUU | GG | UUG |
|  | CamBNRV1 | AUUCUUUUU | GG | UUG |
|  | CanBNRV1 | AUUCUUUUU | GG | UUG |
|  | CarBNRV1 | AUUCUUUUU | GG | UUG |
|  | ChrBNRV1 | AUUCUUUUU | GG | UUG |
|  | CopBNRV1 | AUUCUUUUU | GG | UUG |
|  | DenBNRV1 | AUUCUUUUU | GG | UUG |
|  | DaBNRV1 | AUUCUUUUU | GG | UUG |
|  | EicBNRV1 | AUUCUUUUU | GG | UUG |
|  | LysBNRV1 | AUUCUUUUU | GG | UUG |
|  | MenBNRV1 | AUUCUUUUU | GG | UUG |
|  | NekBNRV1 | AUUCUUUUU | GG | UUG |
|  | NymBNRV1 | AUUCUUUUU | GG | UUG |
|  | OsmBNRV1 | AUUCUUUUU | GG | UUG |
|  | RhoBNRV1 | AUUCUUUUU | GG | UUG |
|  | RicBNRV1 | AUUCUUUUU | GG | UUG |
|  | RorBNRV1 | AUUCUUUUU | GG | UUG |
|  | SalBNRV1 | AUUCUUUUU | GG | UUG |
|  | SalvBNRV1 | AUUCUUUUU | GG | UUG |
|  | StrBNRV1 | AUUCUUUUU | GG | UUG |
|  | ThyBNRV1 | AUUCUUUUU | GG | UUG |
|  | TriBNRV1 | AUUCUUUUU | GG | UUG |
|  | VioBNRV1 | AUUCUUUUU | GG | UUG |
|  | ZizBNRV1 | AUUCUUUUU | GG | UUG |
|  | AaNV | AUUCUUUUU | GG | UUG |
|  | ApVSV | AUUCUUUUU | GG | UUG |
|  | ApRVA | AUUCUUUUU | GG | UUG |
|  | AscSyV2 | AUUCUUUUU | GG | UUG |
|  | BmV2 | AUUCUUUUU | GG | UUG |
|  | BNRV1 | AUUCUUUUU | GG | UUG |
|  | BFTV | AUUCUUUUU | GG | UUG |
|  | BCaRV | AUUCUUUUU | GG | UUG |
|  | CdVCV | AUUCUUUUU | GG | UUG |
|  | CnV1 | AUUCUUUUU | GG | UUG |
|  | DYVV | AUUCUUUUU | GG | UUG |
|  | PyRV1 | AUUCUUUUU | GG | UUG |
|  | PBRV1 | AUUCUUUUU | GG | UUG |
|  | PleArV1 | AUUCUUUUU | GG | UUG |
|  | RhoDeV1 | AUUCUUUUU | GG | UUG |
|  | SaBNV1 | AUUCUUUUU | GG | UUG |
|  | SaBNV2 | AUUCUUUUU | GG | UUG |
|  | SaBNV3 | AUUCUUUUU | GG | UUG |
|  | SaBNV4 | AUUCUUUUU | GG | UUG |
|  | SaBNV5 | AUUCUUUUU | GG | UUG |
|  | SYNV | AUUCUUUUU | GG | UUG |
|  | SYVV | AUUCUUUUU | GG | UUG |
|  | TarBRV1 | AUUCUUUUU | GG | UUG |
|  | TBRV1 | AUUCUUUUU | GG | UUG |
|  | TBRV2 | AUUCUUUUU | GG | UUG |
|  | ZPNRV | AUUCUUUUU | GG | UUG |
| *Deltanucleorhabdovirus* | ArTDNRV1 | AUU(A/C)UUUUU | GAG | UUG |
|  | ArtDNRV2 | AUUAUUUUU | GAG | UUG |
|  | ChrDNRV1 | AUU(A/C)UUUUU | GAG | UUG |
|  | GeuDNRV1 | AUUAUUUUU | GAG | UUG |
|  | MenDNRV1 | AUUCUUUUU | GAG | UUG |
|  | ThyDNRV2 | AUUAUUUUU | GAG | UUG |
|  | TolDNRV1 | AUUAUUUUU | GAG | UUG |
|  | TomDNRV1 | AUU(A/C)UUUUU | GAG | UUG |
|  | VacDNRV1 | AUU(A/C)UUUUU | GAG | UUT |
|  | MSV1 | AUUCUUUUU | GAG | UUG |
|  | StrV3 | AUUCUUUUU | GAG | UUG |
| *Gammanucleorhabdovirus* | BruGNRV1 | AUUUAUUUU | GUAG | UUA |
|  | MyrGNRV1 | AUUUAUUUU | GUAG | UUG |
|  | PopGNRV1 | AUUUAUUUU | GUAG | UU(G/A) |
|  | TamGNRV1 | AUUUAUUUU | GUAG | UUG |
|  | CCMoV | AUUUAUUUU | GUAG | UUG |
|  | MFSV | AUUUAUUUU | GUAG | UUG |

The consensus gene junction sequences of the viruses identified in this study are highlighted in light grey. * Names and abbreviations of newly identified viruses are listed in Tables 1-4; while the names and abbreviations of known viruses are listed in Supp Table 1.
