## Supplementary material for "Expanding the known nucleorhabdovirus world: the final chapter in a trilogy exploring the hidden diversity of plant-associated rhabdoviruses": Table S1

Supplementary Table S1. Names and abbreviations of nucleorhabdoviruses used in this study.

| **Virus name** | **Abbreviation** | **Genus** |
| --- | --- | --- |
| Agave tequilana virus 1 | ATV1 | *Alphanucleorhabdovirus* |
| Artemisia capillaris nucleorhabdovirus 1 | ArtCaNV1 |  |
| babaco nucleorhabdovirus 1 | BabRV1 |  |
| constricta yellow dwarf virus | CYDV |  |
| eggplant mottled dwarf virus | EMDV |  |
| joa yellow blotch associated virus | YYBaV |  |
| maize Iranian mosaic virus | MIMV |  |
| maize mosaic virus | MMV |  |
| Morogoro maize associated virus | MMaV |  |
| peach virus 1 | PeV1 |  |
| potato yellow dwarf virus | PYDV |  |
| Physostegia chlorotic mottle virus | PhCMoV |  |
| rice yellow stunt virus | RYSV |  |
| taro vein chlorosis virus | TVCV |  |
| tomato alphanucleorhabdovirus 1 | TARV1 |  |
| wheat yellow striate virus | WYSV |  |
| Xinjiang alphanucleorhabdovirus | XARV |  |
| alfalfa associated nucleorhabdovirus | AaNV | *Betanucleorhabdovirus* |
| Alpinia vein streaking virus | ApVSV |  |
| apple rootstock virus A | ApRVA |  |
| Asclepias syiriaca virus 2 | AscSyV2 |  |
| Bacopa monnieri virus 2 | BmV2 |  |
| beet betanucleorhabdovirus 1 | BNRV1 |  |
| birds-foot trefoil associated virus | BFTV |  |
| blackcurrant associated rhabdovirus | BCaRV |  |
| cardamom vein clearing virus | CdVCV |  |
| Cnidium virus 1 | CnV1 |  |
| Datura yellow vein virus | DYVV |  |
| Paris yunnanensis rhabdovirus 1 | PyRV1 |  |
| Picris betanucleorhabdovirus 1 | PBRV1 |  |
| Plectranthus aromaticus virus 1 | PleArV1 |  |
| Rhododendron delavayi virus 1 | RhoDeV1 |  |
| Sambucus betanucleorhabdovirus 1 | SaBNV1 |  |
| Sambucus betanucleorhabdovirus 2 | SaBNV2 |  |
| Sambucus betanucleorhabdovirus 3 | SaBNV3 |  |
| Sambucus betanucleorhabdovirus 4 | SaBNV4 |  |
| Sambucus betanucleorhabdovirus 5 | SaBNV5 |  |
| Sonchus yellow net virus | SYNV |  |
| sowthistle yellow vein virus | SYVV |  |
| Taraxacum betanucleorhabdovirus 1 | TarBRV1 |  |
| tomato betanucleorhabdovirus 1 | TBRV1 |  |
| tomato betanucleorhabdovirus 2 | TBRV2 |  |
| Zhuye pepper nucleorhabdovirus | ZPNRV |  |
| Medicago sativa virus 1 | MSV1 | *Deltanucleorhabdovirus* |
| strawberry virus 3 | StrV3 |  |
| cereal chlorotic mottle virus | CCMoV | *Gammanucleorhabdovirus* |
| maize fine streak virus | MFSV |  |
