## Supplementary material for "Expanding the known nucleorhabdovirus world: the final chapter in a trilogy exploring the hidden diversity of plant-associated rhabdoviruses": Table S2

Supplementary Table S2. Nucleotide sequence identity of the complete coding region of alphanucleorhabdoviruses

|  | Aro | ATV1 | ArtCaNV1 | Azo | BabRV1 | Cap | Cas | Cyn | CYDV | EMDV | Hil | JYBaV | Lol | MMV | MIMV | MMaV | PhCMoV | PeV1 | PYDV | Rue | RYSV | Sch | Sen | TaVCV | TARV1 | WYSV | XARV |
| --- | --- | --- | --- | --- | --- | --- | --- | --- | --- | --- | --- | --- | --- | --- | --- | --- | --- | --- | --- | --- | --- | --- | --- | --- | --- | --- | --- |
| AroANRV1 | 100 |  |  |  |  |  |  |  |  |  |  |  |  |  |  |  |  |  |  |  |  |  |  |  |  |  |  |
| ATV1 | 57.4 | 100 |  |  |  |  |  |  |  |  |  |  |  |  |  |  |  |  |  |  |  |  |  |  |  |  |  |
| ArtCaNV1 | 57.9 | 56.9 | 100 |  |  |  |  |  |  |  |  |  |  |  |  |  |  |  |  |  |  |  |  |  |  |  |  |
| AzoANRV1 | 58.9 | 59.1 | 58.9 | 100 |  |  |  |  |  |  |  |  |  |  |  |  |  |  |  |  |  |  |  |  |  |  |  |
| BabRV1 | 57.9 | 57.8 | 58.3 | 58.5 | 100 |  |  |  |  |  |  |  |  |  |  |  |  |  |  |  |  |  |  |  |  |  |  |
| CapANRV1 | 58.7 | 58.7 | 58.9 | 58.8 | 57.9 | 100 |  |  |  |  |  |  |  |  |  |  |  |  |  |  |  |  |  |  |  |  |  |
| CasANRV1 | 58.1 | 58.2 | 57.5 | 58.7 | 58.8 | 58.6 | 100 |  |  |  |  |  |  |  |  |  |  |  |  |  |  |  |  |  |  |  |  |
| CynANRV1 | 56.9 | 57.9 | 57.8 | 57.9 | 57.2 | 58.4 | 58.3 | 100 |  |  |  |  |  |  |  |  |  |  |  |  |  |  |  |  |  |  |  |
| CYDV | 58.2 | 58.6 | 57.9 | 56.6 | 67.7 | 58.4 | 59.2 | 57.5 | 100 |  |  |  |  |  |  |  |  |  |  |  |  |  |  |  |  |  |  |
| EMDV | 58.1 | 58.1 | 58.8 | 58.6 | 61.2 | 58.5 | 58.9 | 58.2 | 60.8 | 100 |  |  |  |  |  |  |  |  |  |  |  |  |  |  |  |  |  |
| HilANRV1 | 57.7 | 58.4 | 59.6 | 58.4 | 57.9 | 59.1 | 58.3 | 57.4 | 57.9 | 58.1 | 100 |  |  |  |  |  |  |  |  |  |  |  |  |  |  |  |  |
| JYBaV | 57.3 | 57.4 | 57.4 | 57.7 | 67.4 | 58.3 | 59.1 | 57.4 | 67.3 | 60.9 | 57.5 | 100 |  |  |  |  |  |  |  |  |  |  |  |  |  |  |  |
| LolANRV1 | 58.5 | 57.3 | 59.4 | 57.6 | 58.6 | 58.5 | 58.3 | 58.5 | 57.9 | 58.8 | 62.6 | 57.8 | 100 |  |  |  |  |  |  |  |  |  |  |  |  |  |  |
| MMV | 57.9 | 57.5 | 58.1 | 58.1 | 57.2 | 57.9 | 57.9 | 63.2 | 58.2 | 58.4 | 57.6 | 57.7 | 58.7 | 100 |  |  |  |  |  |  |  |  |  |  |  |  |  |
| MIMV | 58.4 | 57.4 | 58.1 | 58.9 | 57.7 | 58.3 | 58.1 | 67.1 | 56.9 | 57.3 | 57.6 | 58.1 | 58.2 | 63.1 | 100 |  |  |  |  |  |  |  |  |  |  |  |  |
| MMaV | 58.1 | 57.7 | 58.3 | 58.8 | 56.9 | 57.8 | 58.8 | 63.1 | 58.7 | 58.5 | 57.8 | 57.9 | 58.7 | 63.4 | 63.9 | 100 |  |  |  |  |  |  |  |  |  |  |  |
| PhCMoV | 57.7 | 58.1 | 59.2 | 58.3 | 60.9 | 58.4 | 58.4 | 57.6 | 60.5 | 72.9 | 58.6 | 60.8 | 58.7 | 57.7 | 59.1 | 58.5 | 100 |  |  |  |  |  |  |  |  |  |  |
| PeV1 | 63.1 | 58.2 | 58.5 | 58.1 | 58.3 | 58.3 | 58.2 | 58.3 | 57.2 | 58.4 | 58.3 | 57.6 | 58.7 | 58.1 | 57.9 | 58.5 | 57.6 | 100 |  |  |  |  |  |  |  |  |  |
| PYDV | 58.1 | 57.9 | 57.8 | 58.3 | 66.8 | 57.6 | 58.9 | 57.3 | 67.5 | 60.3 | 58.6 | 67.3 | 58.1 | 58.1 | 58.2 | 59.2 | 60.3 | 58.1 | 100 |  |  |  |  |  |  |  |  |
| RueANRV1 | 57.1 | 57.7 | 57.1 | 56.9 | 58.9 | 58.5 | 59.5 | 58.1 | 59.6 | 58.1 | 57.1 | 58.9 | 58.4 | 58.7 | 57.4 | 58.2 | 58.3 | 57.7 | 59.5 | 100 |  |  |  |  |  |  |  |
| RYSV | 58.2 | 57.3 | 58.9 | 58.8 | 58.6 | 58.8 | 58.4 | 57.1 | 58.1 | 58.6 | 64.1 | 57.9 | 62.5 | 58.3 | 58.4 | 58.9 | 58.6 | 58.1 | 58.9 | 57.2 | 100 |  |  |  |  |  |  |
| SchANRV1 | 57.8 | 58.6 | 58.4 | 58.3 | 57.8 | 60.6 | 58.4 | 57.6 | 58.1 | 58.5 | 58.7 | 57.5 | 58.9 | 57.2 | 58.3 | 57.1 | 58.6 | 58.1 | 58.9 | 58.1 | 58.1 | 100 |  |  |  |  |  |
| SenANRV1 | 57.7 | 58.2 | 57.8 | 58.2 | 57.8 | 58.5 | 58.2 | 58.4 | 57.6 | 57.2 | 57.8 | 57.6 | 57.9 | 58.9 | 58.3 | 57.9 | 57.3 | 58.5 | 58.7 | 58.1 | 58.1 | 57.8 | 100 |  |  |  |  |
| TaVCV | 58.8 | 57.9 | 58.3 | 58.8 | 58.1 | 58.5 | 58.2 | 62.4 | 57.9 | 58.2 | 58.6 | 57.8 | 58.4 | 63.4 | 63.6 | 67.2 | 58.5 | 58.5 | 58.3 | 58.2 | 57.2 | 57.5 | 57.8 | 100 |  |  |  |
| TARV1 | 57.5 | 57.9 | 59.2 | 58.4 | 60.2 | 59.2 | 57.9 | 57.6 | 60.7 | 73.9 | 57.9 | 60.5 | 58.3 | 58.2 | 57.9 | 58.1 | 72.1 | 58.1 | 60.7 | 58.5 | 58.1 | 59.5 | 58.4 | 57.8 | 100 |  |  |
| WYSV | 58.4 | 57.8 | 59.5 | 58.4 | 57.9 | 58.8 | 58.6 | 58.8 | 58.5 | 59.5 | 64.6 | 58.3 | 62.9 | 58.1 | 58.1 | 58.9 | 58.7 | 58.4 | 58.6 | 58.1 | 63.7 | 58.6 | 57.7 | 58.3 | 58.3 | 100 |  |
| XARV | 58.9 | 58.6 | 58.1 | 58.6 | 59.9 | 58.8 | 58.8 | 57.7 | 59.2 | 59.4 | 57.4 | 58.6 | 57.7 | 58.9 | 59.4 | 58.5 | 59.5 | 58.3 | 59.9 | 58.4 | 58.4 | 59.1 | 57.7 | 58.7 | 59.9 | 58.1 | 100 |

* virus names are listed in Supplementary Table S1 and Table 1

Nucleotide sequence identities between 70% and 75% are shaded in violet

Nucleotide sequence identities between 65% and 70% are shaded in orange

Nucleotide sequence identities between 60% and 65% are shaded in green

Nucleotide sequence identities between 55% and 60% are shaded in yellow
