## Supplementary material for "Expanding the known nucleorhabdovirus world: the final chapter in a trilogy exploring the hidden diversity of plant-associated rhabdoviruses": Table S3

Supplementary Table S3. Nucleotide sequence identity of the complete coding region of betanucleorhabdoviruses

|  | AaNV | ApRVA | ApVSV | Art | AscSy  V2 | BFTV | BCaRV | BmV2 | BNRV1 | Boe | Cam | Can | Car | CdVCV | Chr | CnV1 | Cop | Da | Den | DYVV | Eic | Lys | Men | Nek | Nym | Osm |
| --- | --- | --- | --- | --- | --- | --- | --- | --- | --- | --- | --- | --- | --- | --- | --- | --- | --- | --- | --- | --- | --- | --- | --- | --- | --- | --- |
| AaNV | 100 |  |  |  |  |  |  |  |  |  |  |  |  |  |  |  |  |  |  |  |  |  |  |  |  |  |
| ApRVA | 59.8 | 100 |  |  |  |  |  |  |  |  |  |  |  |  |  |  |  |  |  |  |  |  |  |  |  |  |
| ApVSV | 59.5 | 60.3 | 100 |  |  |  |  |  |  |  |  |  |  |  |  |  |  |  |  |  |  |  |  |  |  |  |
| ArtBNRV1 | 59.9 | 60.5 | 60.1 | 100 |  |  |  |  |  |  |  |  |  |  |  |  |  |  |  |  |  |  |  |  |  |  |
| AscSyV2 | 59.8 | 60.4 | 59.7 | 59.8 | 100 |  |  |  |  |  |  |  |  |  |  |  |  |  |  |  |  |  |  |  |  |  |
| BFTV | 58.9 | 59.5 | 60.3 | 59.7 | 59.4 | 100 |  |  |  |  |  |  |  |  |  |  |  |  |  |  |  |  |  |  |  |  |
| BCaRV | 59.7 | 59.6 | 60.9 | 60.2 | 60.7 | 60.7 | 100 |  |  |  |  |  |  |  |  |  |  |  |  |  |  |  |  |  |  |  |
| BmV2 | 59.1 | 59.1 | 60.9 | 60.9 | 59.1 | 61.3 | 59.6 | 100 |  |  |  |  |  |  |  |  |  |  |  |  |  |  |  |  |  |  |
| BNRV1 | 59.3 | 60.1 | 61.2 | 60.7 | 59.9 | 60.4 | 60.8 | 61.3 | 100 |  |  |  |  |  |  |  |  |  |  |  |  |  |  |  |  |  |
| BoeBNRV1 | 59.3 | 58.8 | 60.5 | 60.3 | 58.6 | 60.7 | 60.1 | 61.5 | 64.1 | 100 |  |  |  |  |  |  |  |  |  |  |  |  |  |  |  |  |
| CamBNRV1 | 60.8 | 59.2 | 59.4 | 59.9 | 58.8 | 59.1 | 59.7 | 59.1 | 58.9 | 59.5 | 100 |  |  |  |  |  |  |  |  |  |  |  |  |  |  |  |
| CanBNRV1 | 58.2 | 59.8 | 60.2 | 59.7 | 59.2 | 60.5 | 61.1 | 59.8 | 60.7 | 59.9 | 59.6 | 100 |  |  |  |  |  |  |  |  |  |  |  |  |  |  |
| CarBNRV1 | 58.8 | 59.1 | 59.9 | 59.1 | 59.2 | 61.1 | 60.3 | 61.1 | 63.7 | 64.1 | 58.8 | 58.8 | 100 |  |  |  |  |  |  |  |  |  |  |  |  |  |
| CdVCV | 59.9 | 60.5 | 69.5 | 60.3 | 59.9 | 60.2 | 61.6 | 59.8 | 60.3 | 60.7 | 59.4 | 60.6 | 59.1 | 100 |  |  |  |  |  |  |  |  |  |  |  |  |
| ChrBNRV1 | 59.4 | 60.2 | 59.8 | 64.2 | 60.7 | 60.2 | 61.1 | 60.6 | 60.7 | 59.8 | 59.4 | 59.9 | 59.8 | 60.6 | 100 |  |  |  |  |  |  |  |  |  |  |  |
| CnV1 | 58.7 | 59.4 | 61.4 | 59.9 | 59.5 | 61.1 | 61.1 | 61.4 | 61.6 | 60.7 | 59.4 | 60.8 | 60.4 | 61.7 | 60.7 | 100 |  |  |  |  |  |  |  |  |  |  |
| CopBNRV1 | 59.3 | 59.5 | 60.6 | 59.9 | 59.2 | 62.4 | 60.7 | 62.1 | 62.2 | 60.8 | 58.5 | 60.4 | 61.2 | 60.8 | 60.3 | 59.8 | 100 |  |  |  |  |  |  |  |  |  |
| DaBNRV1 | 58.4 | 59.1 | 60.5 | 59.7 | 57.9 | 61.4 | 60.7 | 60.4 | 60.8 | 60.9 | 59.5 | 59.9 | 61.3 | 60.4 | 60.3 | 60.5 | 61.1 | 100 |  |  |  |  |  |  |  |  |
| DenBNRV1 | 59.3 | 59.6 | 60.7 | 59.9 | 59.5 | 61.3 | 61.2 | 61.3 | 61.3 | 61.2 | 60.1 | 60.5 | 60.9 | 60.6 | 61.4 | 60.6 | 61.8 | 62.7 | 100 |  |  |  |  |  |  |  |
| DYVV | 59.3 | 59.6 | 60.7 | 60.9 | 59.7 | 61.7 | 61.1 | 67.9 | 60.1 | 61.3 | 58.3 | 60.2 | 61.1 | 60.2 | 60.8 | 61.3 | 60.7 | 61.1 | 61.6 | 100 |  |  |  |  |  |  |
| EicBNRV1 | 59.3 | 59.5 | 59.8 | 60.4 | 59.7 | 60.9 | 60.5 | 60.4 | 60.5 | 60.3 | 59.4 | 59.7 | 60.6 | 60.3 | 60.1 | 60.4 | 60.9 | 60.3 | 61.5 | 60.9 | 100 |  |  |  |  |  |
| LysBNRV1 | 60.3 | 63.6 | 61.1 | 60.5 | 61.4 | 60.5 | 60.1 | 60.3 | 60.9 | 59.1 | 59.9 | 60.6 | 58.2 | 60.7 | 60.6 | 60.1 | 59.6 | 59.7 | 60.6 | 59.8 | 60.5 | 100 |  |  |  |  |
| MenBNRV1 | 59.5 | 59.2 | 61.9 | 60.7 | 59.3 | 60.6 | 60.4 | 60.1 | 60.1 | 60.3 | 59.3 | 60.4 | 59.8 | 61.7 | 60.9 | 61.9 | 60.6 | 60.8 | 60.9 | 60.5 | 60.7 | 59.4 | 100 |  |  |  |
| NekBNRV1 | 61.2 | 60.4 | 58.5 | 59.6 | 60.4 | 58.8 | 58.7 | 59.2 | 59.7 | 58.4 | 65.7 | 58.8 | 59.1 | 59.3 | 60.3 | 58.4 | 58.8 | 58.8 | 60.1 | 58.8 | 59.1 | 60.7 | 59.2 | 100 |  |  |
| NymBNRV1 | 58.5 | 59.3 | 60.1 | 60.7 | 60.3 | 60.9 | 60.5 | 60.1 | 60.6 | 60.9 | 59.3 | 60.3 | 60.4 | 59.4 | 59.9 | 60.6 | 61.9 | 60.1 | 60.8 | 61.4 | 65.2 | 60.2 | 60.8 | 58.7 | 100 |  |
| OsmBNRV1 | 59.4 | 59.5 | 60.4 | 60.1 | 60.1 | 61.2 | 59.8 | 61.5 | 61.4 | 61.8 | 58.7 | 59.7 | 61.7 | 60.8 | 60.2 | 60.7 | 62.6 | 61.3 | 62.4 | 60.7 | 60.8 | 60.3 | 60.5 | 58.9 | 60.8 | 100 |
| PBRV1 | 59.8 | 60.8 | 61.3 | 63.1 | 61.6 | 60.5 | 60.1 | 60.3 | 60.2 | 60.9 | 59.7 | 59.9 | 59.9 | 61.3 | 63.2 | 60.9 | 60.6 | 58.7 | 61.2 | 59.9 | 60.4 | 60.3 | 60.7 | 59.1 | 60.4 | 60.1 |
| PyRV1 | 59.5 | 58.7 | 60.9 | 60.1 | 59.4 | 61.3 | 60.7 | 60.8 | 60.2 | 59.9 | 59.6 | 60.5 | 60.6 | 60.9 | 61.1 | 60.5 | 59.9 | 60.1 | 60.7 | 60.9 | 60.9 | 59.4 | 60.9 | 59.3 | 60.5 | 60.6 |
| PleArV1 | 59.8 | 59.5 | 61.1 | 60.1 | 59.9 | 61.6 | 60.9 | 64.7 | 61.4 | 61.7 | 59.2 | 59.9 | 61.6 | 60.1 | 60.6 | 60.4 | 61.4 | 61.2 | 61.3 | 65.3 | 60.7 | 59.5 | 60.9 | 58.7 | 60.5 | 61.3 |
| RhoBNRV1 | 59.6 | 59.8 | 61.1 | 60.9 | 59-9 | 59.5 | 60.7 | 60.3 | 60.3 | 59.9 | 59.4 | 60.1 | 59.8 | 60.2 | 60.8 | 59.3 | 59.9 | 60.3 | 60.8 | 59.5 | 60.1 | 60.5 | 60.2 | 59.5 | 59.5 | 59.9 |
| RhoDeV1 | 59.3 | 59.9 | 60.7 | 61.1 | 60.9 | 60.5 | 60.5 | 59.9 | 59.8 | 59.8 | 60.2 | 59.9 | 59.8 | 60.1 | 61.1 | 60.9 | 59.4 | 59.1 | 59.9 | 59.8 | 60.7 | 60.3 | 59.8 | 58.9 | 60.3 | 60.1 |
| RicBNRV1 | 59.4 | 59.7 | 61.2 | 60.1 | 60.2 | 60.9 | 61.7 | 60.9 | 61.8 | 60.9 | 58.9 | 60.2 | 60.1 | 60.1 | 60.7 | 61.2 | 61.1 | 61.9 | 62.9 | 60.9 | 61.1 | 60.6 | 60.6 | 58.9 | 60.1 | 61.8 |
| RorBNRV1 | 59.6 | 60.3 | 60.9 | 60.8 | 59.8 | 62.1 | 61.5 | 61.2 | 61.6 | 61.6 | 59.2 | 61.6 | 60.8 | 60.9 | 60.7 | 60.5 | 62.6 | 60.8 | 61.5 | 61.5 | 61.5 | 60.6 | 60.1 | 59.8 | 60.9 | 61.9 |
| SaBNV1 | 59.1 | 59.4 | 60.5 | 59.4 | 59.1 | 60.8 | 60.2 | 61.1 | 67.3 | 63.2 | 58.5 | 59.6 | 62.8 | 60.4 | 58.9 | 60.6 | 61.5 | 60.4 | 60.1 | 61.3 | 60.4 | 59.6 | 60.2 | 58.9 | 60.6 | 61.3 |
| SaBNV2 | 58.5 | 58.3 | 60.2 | 58.9 | 59.8 | 60.6 | 60.2 | 61.3 | 66.9 | 63.9 | 58.7 | 60.1 | 63.9 | 59.7 | 58.7 | 61.8 | 60.9 | 60.6 | 60.5 | 61.8 | 60.2 | 59.4 | 60.4 | 58.5 | 59.8 | 61.2 |
| SaBNV3 | 59.3 | 58.8 | 59.9 | 59.8 | 59.3 | 60.9 | 60.5 | 61.3 | 67.1 | 63.4 | 58.9 | 59.9 | 62.7 | 59.9 | 59.7 | 60.4 | 61.6 | 60.3 | 60.8 | 61.8 | 60.2 | 58.6 | 60.2 | 58.8 | 60.6 | 61.5 |
| SaBNV4 | 59.3 | 58.6 | 59.8 | 59.4 | 59.7 | 60.4 | 60.4 | 61.6 | 67.5 | 64.3 | 57.9 | 60.1 | 63.6 | 59.8 | 59.3 | 60.9 | 61.5 | 60.7 | 61.3 | 61.6 | 60.7 | 59.8 | 60.1 | 58.5 | 60.2 | 61.6 |
| SaBNV5 | 58.7 | 57.7 | 60.2 | 59.8 | 58.8 | 60.2 | 60.8 | 61.8 | 66.9 | 63.7 | 58.4 | 59.6 | 63.3 | 59.9 | 59.5 | 60.9 | 60.9 | 60.3 | 60.6 | 61.2 | 60.2 | 60.2 | 60.3 | 58.4 | 60.1 | 61.5 |
| SalBNRV1 | 59.3 | 59.9 | 60.6 | 60.2 | 60.2 | 59.4 | 60.5 | 59.6 | 60.9 | 58.8 | 58.9 | 58.9 | 59.2 | 59.6 | 60.3 | 58.9 | 58.8 | 59.2 | 59.8 | 59.9 | 59.9 | 60.2 | 58.9 | 59.1 | 59.8 | 59.4 |
| SalvBNRV1 | 59.6 | 59.9 | 60.2 | 60.1 | 59.1 | 64.3 | 60.7 | 61.2 | 61.2 | 61.1 | 59.6 | 60.6 | 61.1 | 60.8 | 59.9 | 61.2 | 62.8 | 59.9 | 61.9 | 61.3 | 61.1 | 60.1 | 60.7 | 59.3 | 60.8 | 62.1 |
| StrBNRV1 | 59.2 | 60.3 | 60.2 | 60.6 | 60.1 | 60.8 | 60.7 | 60.4 | 60.7 | 60.1 | 59.6 | 60.5 | 59.8 | 60.6 | 60.5 | 60.3 | 60.5 | 60.7 | 61.5 | 61.1 | 70.8 | 60.2 | 60.7 | 59.7 | 64.9 | 60.8 |
| SYNV | 59.2 | 59.1 | 60.6 | 60.7 | 60.4 | 60.2 | 60.7 | 59.9 | 60.2 | 60.5 | 59.4 | 59.9 | 59.9 | 61.2 | 60.1 | 60.7 | 60.3 | 60.1 | 59.9 | 60.2 | 60.1 | 60.3 | 61.5 | 59.1 | 60.1 | 59.5 |
| SYVV | 58.6 | 59.1 | 60.9 | 60.3 | 59.2 | 60.9 | 61.8 | 60.6 | 61.2 | 61.1 | 59.9 | 59.8 | 60.9 | 60.8 | 60.7 | 60.6 | 61.7 | 61.1 | 61.8 | 61.1 | 60.3 | 60.1 | 60.3 | 59.1 | 59.6 | 61.3 |
| TarBRV1 | 59.1 | 59.4 | 61.3 | 59.2 | 59.3 | 59.6 | 60.2 | 60.1 | 59.9 | 59.5 | 58.5 | 59.3 | 59.7 | 60.1 | 59.2 | 60.2 | 59.6 | 59.7 | 60.2 | 60.1 | 60.4 | 59.6 | 59.1 | 59.2 | 60.1 | 59.8 |
| TBRV1 | 59.1 | 57.6 | 60.5 | 60.2 | 59.1 | 60.5 | 60.4 | 60.6 | 67.2 | 63.7 | 58.1 | 60.4 | 63.6 | 59.9 | 59.2 | 60.9 | 61.4 | 60.2 | 60.8 | 61.7 | 60.8 | 59.1 | 60.6 | 59.1 | 59.2 | 61.1 |
| TBRV2 | 59.4 | 59.2 | 60.8 | 59.7 | 58.7 | 61.2 | 61.2 | 61.7 | 68.3 | 63.7 | 58.8 | 60.6 | 64.5 | 60.2 | 60.1 | 60.5 | 61.2 | 59.8 | 61.2 | 61.7 | 60.9 | 59.2 | 60.4 | 58.8 | 60.2 | 61.3 |
| ThyBNRV1 | 59.1 | 59.3 | 60.9 | 60.6 | 59.9 | 64.5 | 61.1 | 61.2 | 61.8 | 61.4 | 59.4 | 60.8 | 61.3 | 61.3 | 60.5 | 60.9 | 62.2 | 60.3 | 61.1 | 60.8 | 61.2 | 60.2 | 60.8 | 59.3 | 61.2 | 62.7 |
| TriBNRV1 | 58.5 | 58.8 | 60.3 | 60.2 | 59.7 | 60.7 | 60.3 | 61.1 | 63.1 | 63.3 | 59.6 | 59.6 | 63.3 | 60.7 | 60.4 | 60.2 | 61.1 | 60.2 | 60.1 | 60.6 | 61.1 | 60.1 | 60.1 | 59.5 | 60.3 | 61.1 |
| VioBNRV1 | 59.5 | 59.8 | 60.6 | 60.5 | 59.2 | 60.4 | 61.6 | 60.5 | 60.5 | 60.3 | 59.8 | 60.9 | 59.5 | 60.2 | 60.8 | 61.1 | 59.2 | 60.3 | 60.7 | 60.9 | 61.2 | 61.1 | 59.9 | 59.4 | 60.7 | 59.6 |
| ZizBNRV1 | 59.3 | 59.6 | 61.1 | 60.8 | 60.4 | 65.1 | 60.7 | 60.9 | 61.8 | 61.3 | 59.3 | 60.1 | 60.9 | 60.5 | 60.9 | 61.1 | 62.1 | 60.3 | 61.7 | 61.4 | 61.3 | 60.3 | 60.8 | 58.9 | 61.1 | 62.4 |
| ZPNRV | 56.8 | 57.4 | 56.6 | 57.3 | 58.1 | 57.8 | 57.1 | 56.4 | 58.2 | 57.6 | 57.6 | 57.8 | 56.3 | 55.9 | 57.4 | 57.6 | 56.9 | 56.9 | 56.7 | 56.7 | 56.9 | 57.9 | 55.5 | 58.1 | 57.5 | 56.5 |
|  | PBRV1 | PyRV1 | PleAr  V1 | Rho | RhoDeV1 | Ric | Ror | SaBNV1 | SaBNV2 | SaBNV3 | SaBNV4 | SaBNV5 | Sal | Salv | Str | SYNV | SYVV | TarBRV1 | TBRV1 | TBRV2 | Thy | Tri | Vio | Ziz | ZPNRV |  |
| PBRV1 | 100 |  |  |  |  |  |  |  |  |  |  |  |  |  |  |  |  |  |  |  |  |  |  |  |  |  |
| PyRV1 | 61.2 | 100 |  |  |  |  |  |  |  |  |  |  |  |  |  |  |  |  |  |  |  |  |  |  |  |  |
| PleArV1 | 60.2 | 60.4 | 100 |  |  |  |  |  |  |  |  |  |  |  |  |  |  |  |  |  |  |  |  |  |  |  |
| RhoBNRV1 | 60.3 | 60.2 | 59.5 | 100 |  |  |  |  |  |  |  |  |  |  |  |  |  |  |  |  |  |  |  |  |  |  |
| RhoDeV1 | 60.9 | 60.3 | 60.5 | 65.8 | 100 |  |  |  |  |  |  |  |  |  |  |  |  |  |  |  |  |  |  |  |  |  |
| RicBNRV1 | 61.7 | 61.2 | 60.8 | 59.7 | 60.4 | 100 |  |  |  |  |  |  |  |  |  |  |  |  |  |  |  |  |  |  |  |  |
| RorBNRV1 | 60.5 | 61.4 | 62.3 | 61.1 | 60.5 | 61.4 | 100 |  |  |  |  |  |  |  |  |  |  |  |  |  |  |  |  |  |  |  |
| SaBNV1 | 59.6 | 59.4 | 60.9 | 60.1 | 59.9 | 60.6 | 60.6 | 100 |  |  |  |  |  |  |  |  |  |  |  |  |  |  |  |  |  |  |
| SaBNV2 | 59.8 | 60.3 | 61.1 | 59.9 | 59.1 | 60.9 | 61.7 | 71.5 | 100 |  |  |  |  |  |  |  |  |  |  |  |  |  |  |  |  |  |
| SaBNV3 | 59.8 | 59.9 | 61.1 | 60.1 | 59.7 | 61.5 | 61.5 | 71.9 | 73.1 | 100 |  |  |  |  |  |  |  |  |  |  |  |  |  |  |  |  |
| SaBNV4 | 59.5 | 59.8 | 60.8 | 59.8 | 59.7 | 60.9 | 61.4 | 71.4 | 72.7 | 74.1 | 100 |  |  |  |  |  |  |  |  |  |  |  |  |  |  |  |
| SaBNV5 | 60.4 | 60.2 | 60.9 | 59.8 | 59.1 | 60.9 | 60.9 | 72.4 | 73.9 | 74.8 | 74.9 | 100 |  |  |  |  |  |  |  |  |  |  |  |  |  |  |
| SalBNRV1 | 60.9 | 59.9 | 59.4 | 60.6 | 61.2 | 59.1 | 59.8 | 59.3 | 59.7 | 59.5 | 59.6 | 59.4 | 100 |  |  |  |  |  |  |  |  |  |  |  |  |  |
| SalvBNRV1 | 60.2 | 61.1 | 61.6 | 59.8 | 60.1 | 61.7 | 61.7 | 60.8 | 61.1 | 60.8 | 60.8 | 61.4 | 59.9 | 100 |  |  |  |  |  |  |  |  |  |  |  |  |
| StrBNRV1 | 60.8 | 60.4 | 60.8 | 60.9 | 60.1 | 61.2 | 61.1 | 60.4 | 59.9 | 60.1 | 61.2 | 60.7 | 60.4 | 61.2 | 100 |  |  |  |  |  |  |  |  |  |  |  |
| SYNV | 61.1 | 61.8 | 60.9 | 60.1 | 60.3 | 60.7 | 61.1 | 59.9 | 59.8 | 60.1 | 60.8 | 60.2 | 59.5 | 60.4 | 59.9 | 100 |  |  |  |  |  |  |  |  |  |  |
| SYVV | 60.4 | 60.6 | 61.4 | 59.6 | 59.3 | 61.2 | 61.5 | 61.1 | 60.4 | 60.3 | 60.7 | 59.9 | 59.3 | 61.6 | 60.3 | 59.6 | 100 |  |  |  |  |  |  |  |  |  |
| TarBRV1 | 60.2 | 61.7 | 59.9 | 59.9 | 59.7 | 59.7 | 60.6 | 59.4 | 60.1 | 59.8 | 60.2 | 59.2 | 59.5 | 59.7 | 60.5 | 61.5 | 59.2 | 100 |  |  |  |  |  |  |  |  |
| TBRV1 | 59.9 | 59.6 | 61.3 | 59.8 | 59.8 | 60.6 | 61.1 | 68.8 | 66.9 | 67.7 | 67.8 | 68.3 | 59.5 | 61.3 | 59.9 | 60.4 | 59.6 | 59.5 | 100 |  |  |  |  |  |  |  |
| TBRV2 | 59.9 | 60.4 | 61.4 | 59.8 | 59.9 | 61.2 | 61.7 | 68.8 | 68.1 | 67.6 | 68.2 | 68.2 | 59.4 | 61.3 | 60.4 | 59.7 | 60.7 | 60.3 | 68.1 | 100 |  |  |  |  |  |  |
| ThyBNRV1 | 60.3 | 60.2 | 61.6 | 60.7 | 60.1 | 61.4 | 61.7 | 60.8 | 61.3 | 61.2 | 60.8 | 60.8 | 59.3 | 71.9 | 61.8 | 60.3 | 61.3 | 60.8 | 60.7 | 61.3 | 100 |  |  |  |  |  |
| TriBNRV1 | 59.8 | 60.5 | 60.6 | 59.1 | 60.4 | 60.1 | 60.1 | 63.2 | 62.5 | 62.6 | 62.9 | 63.1 | 59.3 | 60.9 | 60.7 | 60.2 | 59.8 | 59.8 | 63.4 | 63.7 | 61.1 | 100 |  |  |  |  |
| VioBNRV1 | 60.7 | 68.3 | 60.9 | 60.7 | 60.4 | 60.6 | 60.2 | 59.9 | 59.5 | 59.7 | 60.1 | 59.7 | 59.6 | 60.7 | 61.3 | 62.2 | 59.8 | 61.9 | 59.9 | 59.6 | 60.5 | 59.9 | 100 |  |  |  |
| ZizBNRV1 | 60.2 | 60.2 | 61.3 | 60.3 | 60.2 | 61.6 | 61.8 | 61.3 | 60.8 | 61.1 | 61.9 | 61.3 | 59.6 | 71.4 | 60.9 | 59.9 | 61.3 | 60.1 | 61.3 | 61.1 | 74.1 | 60.6 | 60.9 | 100 |  |  |
| ZPNRV | 57.6 | 57.6 | 56.6 | 56.6 | 57.6 | 57.1 | 57.1 | 57.9 | 56.9 | 55.8 | 56.6 | 56.8 | 56.9 | 57.9 | 57.1 | 57.1 | 56.9 | 57.3 | 57.2 | 56.7 | 57.1 | 56.8 | 57.2 | 56.7 | 100 |  |

* virus names are listed in Supplementary Table S1 and Table 2

Nucleotide sequence identities between 70% and 75% are shaded in violet

Nucleotide sequence identities between 65% and 70% are shaded in orange

Nucleotide sequence identities between 60% and 65% are shaded in green

Nucleotide sequence identities between 55% and 60% are shaded in yellow
