## Supplementary material for "Expanding the known nucleorhabdovirus world: the final chapter in a trilogy exploring the hidden diversity of plant-associated rhabdoviruses": Table S4

Supplementary Table S4. Nucleotide sequence identity of the complete coding region of gammanucleorhabdoviruses

|  | BruGNRV1 | CCMoV | MFSV | MyrGNRV1 | PopGNRV1 | TamGNRV1 |
| --- | --- | --- | --- | --- | --- | --- |
| BruGNRV1 | 100 |  |  |  |  |  |
| CCMoV | 60.1 | 100 |  |  |  |  |
| MFSV | 59.8 | 63.3 | 100 |  |  |  |
| MyrGNRV1 | 60.9 | 59.3 | 59.3 | 100 |  |  |
| PopGNRV1 | 61.9 | 58.7 | 59.4 | 60.7 | 100 |  |
| TamGNRV1 | 59.4 | 59.7 | 59.7 | 58.9 | 60.1 | 100 |

* virus names are listed in Supplementary Table S1 and Table 3

Nucleotide sequence identities between 60% and 65% are shaded in green

Nucleotide sequence identities between 55% and 60% are shaded in yellow
