## Supplementary material for "Expanding the known nucleorhabdovirus world: the final chapter in a trilogy exploring the hidden diversity of plant-associated rhabdoviruses": Table S5

Supplementary Table S5. Nucleotide sequence identity of the complete coding region of deltanucleorhabdoviruses

|  | ArtDNRV1 | ChrDNRV1 | GeuDNRV1 | MenDNRV1 | MSV1 | StrV3 | ThyDNRV1 | TolDNRV1 | TomDNRV1 | VacDNRV1 |
| --- | --- | --- | --- | --- | --- | --- | --- | --- | --- | --- |
| ArtDNRV1 | 100 |  |  |  |  |  |  |  |  |  |
| ChrDNRV1 | 66.9 | 100 |  |  |  |  |  |  |  |  |
| GeuDNRV1 | 59.9 | 59.2 | 100 |  |  |  |  |  |  |  |
| MenDNRV1 | 63.4 | 62.7 | 58.3 | 100 |  |  |  |  |  |  |
| MSV1 | 62.6 | 62.9 | 58.8 | 64.6 | 100 |  |  |  |  |  |
| StrV3 | 62.9 | 62.5 | 58.5 | 65.8 | 64.7 | 100 |  |  |  |  |
| ThyDNRV1 | 62.8 | 62.3 | 58.8 | 64.9 | 64.9 | 68.5 | 100 |  |  |  |
| TolDNRV1 | 62.2 | 61.7 | 59.3 | 62.1 | 62.3 | 62.7 | 62.8 | 100 |  |  |
| TomDNRV1 | 62.1 | 62.2 | 59.1 | 63.6 | 64.9 | 63.8 | 64.1 | 61.9 | 100 |  |
| VacDNRV1 | 62.8 | 62.3 | 58.9 | 64.9 | 64.3 | 70.7 | 68.2 | 61.8 | 63.5 | 100 |

* virus names are listed in Supplementary Table S1 and Table 4

Nucleotide sequence identities between 70% and 75% are shaded in violet

Nucleotide sequence identities between 65% and 70% are shaded in orange

Nucleotide sequence identities between 60% and 65% are shaded in green

Nucleotide sequence identities between 55% and 60% are shaded in yellow
